# AMPK-dependent FLCN phosphorylation activates a survival response to hypoxia

**DOI:** 10.64898/2026.05.20.726614

**Authors:** Josue M. J. Ramirez-Reyes, Ariadna Pollato-Blanco, Stephanie Duhamel, Steven Hébert, Claudia L. Kleinman, Bhushan Nagar, Morag Park, Josie Ursini-Siegel, Arnim Pause

**Affiliations:** Goodman Cancer Research Institute, McGill University; Montreal, Quebec, Canada; Department of Biochemistry, McGill University; Montreal, Quebec, Canada; Lady Davis Institute for Medical Research, Jewish General Hospital; Montreal, Quebec, Canada; Department of Human Genetics, McGill University; Montreal, Quebec, Canada; Division of Experimental Medicine, McGill University; Montreal, Quebec, Canada; Department of Medicine, McGill University; Montreal, Quebec, Canada; Department of Oncology, McGill University; Montreal, Quebec, Canada

**Keywords:** FLCN, AMPK, mTORC1, TFEB, TFE3, Hypoxia

## Abstract

Adaptation to low oxygen tensions is required to survive physiological and pathological stress conditions. We discovered an adenosine monophosphate–activated protein kinase (AMPK)-dependent, hypoxia-specific survival pathway through transcription factors (TFEB, TFE3). This pathway is distinct from nutrient sensing and independent of GAP Activity Towards Rags 1 (GATOR1). Mechanistically, AMPK phosphorylates the tumor suppressor folliculin (FLCN) at a highly conserved serine, inhibiting its GAP activity towards the Rag-GTPases, resulting in mechanistic target of rapamycin complex 1 (mTORC1) inhibition, TFE3 activation, resulting in lysosomal and mitochondrial biogenesis essential for hypoxia adaptation. TFEB/3 activity correlates with the hypoxia signature in hypoxic tumor areas. Cancer patients with elevated TFEB/3 activity show poor survival, specifically in hypoxia. These findings identify the AMPK-FLCN-TFE3 axis as a therapeutic target in tumors.

## Introduction

Hypoxia, a reduction in oxygen availability, occurs in physiological contexts as well as in pathological conditions such as stroke, myocardial infarction, and cancer (Kumar and Choi, 2015). Cells experiencing hypoxia face dual challenges: reduced ATP production by impaired oxidative phosphorylation, accompanied by reactive oxygen species (ROS) accumulation that causes cellular damage (Zhao et al., 2020). Although the hypoxia-inducible factor 1-alpha (HIF-1α), the mechanistic Target Of Rapamycin Complex 1 mTORC1, and the AMP-activated protein kinase (AMPK) sit as the central mechanisms in response to hypoxia (Xηυν ανδ κιμ, 2021; Σεμενζα, 2001), how these signaling networks are integrated remains incompletely understood.

mTORC1 and AMPK are the two major metabolic hubs that coordinate anabolic and catabolic responses to energetic and redox stress (Gonzalez et al., 2020). While hypoxia modulates mTORC1-mediated protein synthesis via REDD1, ATF, and PERK, hypoxia also activates AMPK via mitochondrial ROS-dependent CaMKKβ activation (Brugarolas et al., 2004; Harding et al., 2003; Koumenis et al., 2002; Mungai et al., 2011). Although AMPK antagonizes mTORC1 during energetic stress, how hypoxia links AMPK to mTORC1 remains poorly defined.

Canonically, AMPK inhibits mTORC1 through phosphorylating TSC2 and/or Raptor, primarily affecting eukaryotic elongation factor 2 (eEF2), S6 kinases and 4E-BPs, implicated in the regulation of protein synthesis. In this way, AMPK inhibits mTORC1-mediated translation, (Gwinn et al., 2008; Inoki et al., 2003). However, mTORC1 and AMPK catabolic targets include the transcription factors TFEB and TFE3 (TFEB/3) involved in autophagy and lysosomal biogenesis (Martina et al., 2012; Paquette et al., 2021).

TFEB/3 have emerged as master regulators of cellular stress responses, coordinating expression of lysosomal, autophagic, and mitochondrial genes (El-Houjeiri et al., 2019; Erlich et al., 2018; Malik et al., 2023; Martina et al., 2014; Martina and Puertollano, 2018; Settembre et al., 2013). Their activity is primarily controlled by mTORC1, which phosphorylates TFEB/3 to retain them in the cytoplasm in a Rheb and growth factors-independent manner, making TFEB/3 a different mTORC1 functional branch (Martina *et al*., 2012; Settembre et al., 2012). When mTORC1 is inhibited, TFEB/3 translocate to the nucleus and activate transcriptional programs that enhance autophagy together with lysosomal and mitochondrial biogenesis (Erlich *et al*., 2018; Malik *et al*., 2023; Martina *et al*., 2014; Settembre *et al*., 2013). This regulatory axis is activated during amino acid starvation, in which mTORC1 is inactivated through the inactivation of the Rag-GTPases heterodimeric complexes, composed of RagA/B with RagC/D subunits, which act as molecular switches controlling mTORC1 recruitment to lysosomes (Cui et al., 2023; Gollwitzer et al., 2022; Martina and Puertollano, 2013; Napolitano et al., 2020).

Two GTPase-activating protein (GAP) complexes regulate Rag-GTPases nucleotide states: GAP Activity Toward RAGs (GATOR1) acts on RagA/B, while folliculin (FLCN) in complex with Folliculin Interacting Protein 1 or 2 (FNIP1/2) acts on RagC/D (Bar-Peled et al., 2013; Tsun et al., 2013). FLCN is a tumor suppressor gene responsible for the Birt-Hogg-Dube (BHD) syndrome, an autosomal dominant genetic condition characterized by lung cysts, skin fibrofolliculomas and increased risk for developing bilateral renal tumors (Schmidt and Linehan, 2018). BHD patients develop kidney tumors specifically in the chronically hypoxic renal medulla, suggesting a connection between FLCN function and oxygen homeostasis that has not been molecularly defined (Brezis and Rosen, 1995; Liu et al., 2022).

Pharmacological AMPK activation induces TFEB/3 nuclear translocation (Malik *et al*., 2023; Paquette *et al*., 2021). However, whether AMPK physiologically regulates TFEB/3 in response to specific stresses, and the mechanism underlying this regulation, remain unknown. Interaction of AMPK with the FLCN/FNIP GAP complex was reported but no functional consequences were described (Baba et al., 2006; Hasumi et al., 2008; Takagi et al., 2008). Recent work demonstrated that pharmacological AMPK activation induces FNIP1 phosphorylation at five serine residues, proposing that this regulates lysosomal and mitochondrial biogenesis (Malik *et al*., 2023). However, this study used pharmacological AMPK activators and did not identify phosphorylation sites in FLCN, FNIP2, or changes in GAP activity, leaving critical mechanistic questions unanswered.

We previously demonstrated that FLCN regulates AMPK-dependent autophagy and adaptation to metabolic stress, including heat, anoxia, and oxidative stress (El-Houjeiri et al., 2021; El-Houjeiri *et al*., 2019; Possik et al., 2015; Possik et al., 2014; Yan et al., 2016; Yan et al., 2014). Furthermore, we showed that FLCN loss promotes AMPK-dependent metabolic rewiring, including activation of aerobic glycolysis, oxidative phosphorylation and HIF-1α-dependent programs (Possik *et al*., 2014; Yan *et al*., 2014), and that the FLCN-AMPK axis regulates innate immune responses and pathogen resistance through TFEB/3 (El-Houjeiri *et al*., 2019).

Furthermore, we reported that loss of FLCN in luminal breast cancer cells prompts metabolic reprogramming that increases cellular bioenergetics and angiogenesis. Importantly, these alterations that increase metabolic fitness of FLCN-deficient cells are driven through TFE3 transcriptional activation (El-Houjeiri *et al*., 2021). Therefore, FLCN emerges as a critical factor regulating mTORC1 and AMPK, which govern TFEB/3 activity. However, the mechanisms governing FLCN regulation and energy/nutrient-state sensing have remained elusive for more than a decade. Moreover, the functional interplay between AMPK and FLCN, and the extent to which they regulate one another, remains unresolved. Therefore, the molecular mechanisms of the AMPK-FLCN-mTORC1-TFEB/3 axis and the physiological regulator remain uncovered.

Here, we report that hypoxia serves as a physiological stimulus activating the AMPK-FLCN-mTORC1-TFEB/3 pathway through a mechanism distinct from amino acid and glucose sensing. We identify AMPK-dependent FLCN phosphorylation at the highly conserved serine 130 as the key regulatory event. This modification inhibits GAP activity of FLCN, preventing Rag GTPase activation and thereby releasing TFEB/3 from mTORC1 control. Importantly, this pathway operates independently of the amino acid-responsive GAP GATOR1, revealing that oxygen-dependent regulation and amino acids are sensed via distinctive pathways. This hypoxia-dependent and nutrient-independent adaptation pathway uncovers an AMPK-dependent mTORC1 regulation mechanism under hypoxia via FLCN, resulting in TFEB/3 activation, which is a previously unappreciated mechanism of hypoxia adaptation. This pathway provides extra plasticity and flexibility to hypoxia in addition to other adaptation pathways such as HIF-1α, ATF4, NF-κB and p53 (Culver et al., 2010; Harding *et al*., 2003; Semenza, 2001; Zhang et al., 2021). In human tumors, where hypoxia is a common microenvironmental feature, the HIF-1α-dependent hypoxia signature correlated with increased TFE3 activity. In addition, high TFEB/3 activity in hypoxic but not normoxic tumors is associated with poor prognosis. Our findings reveal a dedicated oxygen-sensing mechanism with important implications for human physiology and cancer biology, demonstrating the mechanistic and physiological function of the AMPK-FLCN-mTORC1-TFEB/3 axis (El-Houjeiri *et al*., 2021; El-Houjeiri *et al*., 2019; Paquette *et al*., 2021; Possik *et al*., 2015; Possik *et al*., 2014; Yan *et al*., 2016; Yan *et al*., 2014).

## Results

### Hypoxia induces AMPK-dependent TFE3 nuclear translocation and transcriptional activity

Building on our previous finding that AMPK-dependent phosphorylation is required for TFEB/3 transcriptional activation (Paquette *et al*., 2021), we investigated whether hypoxia (a physiological AMPK activator) regulates TFEB/3. Our prior work in *C. elegans* demonstrated that loss of FLCN confers AMPK-dependent resistance to metabolic stresses, including oxidative stress (paraquat, H_2_O_2_), heat, and anoxia, stresses that all impair mitochondrial function and severely deplete cellular energy (Possik *et al*., 2014). We exposed wild-type (WT) and AMPK-deficient (AMPK DKO) mouse embryonic fibroblasts (MEFs) to 1% O_2_ for 2 hours. Immunofluorescence revealed that TFE3 translocated to the nucleus in WT cells under hypoxia, but AMPK DKO cells showed enhanced cytoplasmic retention of TFE3 during hypoxia compared to normoxia (Fig. 1A, B), suggesting hyperactivation of the mTORC1-TFEB/3 branch. In stark contrast, amino acid starvation (EBSS) induced TFE3 nuclear translocation equally in both WT and AMPK DKO cells (Fig. 1A, B). Glucose starvation also induced TFE3 translocation in an AMPK-independent manner, demonstrating that AMPK is specifically required for hypoxia-induced TFE3 nuclear translocation, unlike amino acid and glucose sensing stress responses (fig. S1A, B). This hypoxia response was confirmed in human HEK293T cells (fig. S1C-E).

**Fig. 1.**
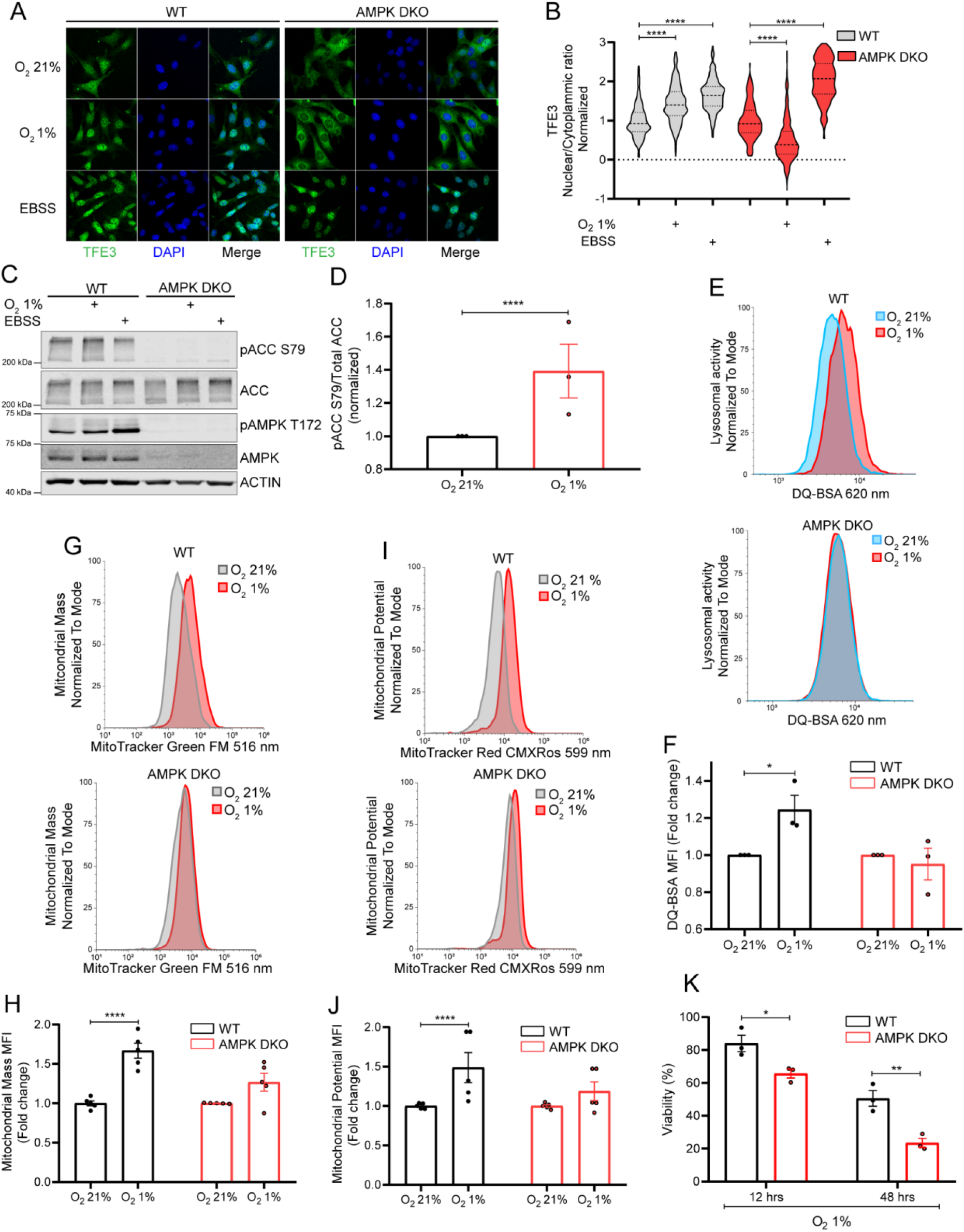
TFE3 nuclear translocation requires AMPK in hypoxia but not starvation and induces cell survival. **(A)** Representative immunofluorescence images of TFE3 localization in the indicated cell lines and conditions. (B) Nuclear/cytoplasmic ratio quantification of A. Data are normalized to the control condition of each cell line. (C) Immunoblotting of AMPK activation in normal, starvation or hypoxic conditions in WT and AMPK DKO MEFs. (D) Phospho-ratio quantification of ACC from C. (E) Representative histograms of lysosomal activity measured by DQ-BSA by flow cytometry in the indicated cell lines incubated in hypoxia for two hours. (F) Quantification of the mean fluorescent intensity of E. Data are normalized to the control condition of each cell line. (G) Representative histograms of mitochondrial mass of the indicated cell lines incubated in hypoxia for 24 hours. (H) Quantification of the mean fluorescent intensity of G. (I) Representative histograms of mitochondrial potential of the indicated cell lines incubated in hypoxia for 24 hours. (J) Quantification of the mean fluorescent intensity of I. (K) Cell viability in the indicated cell lines incubated in hypoxia for 12 hours or 48 hours. Data are represented as the mean ± SEM. A minimum of three independent experiments were performed, *p-value < 0.05, **p-value < 0.01, **** p-value < 0.0001.

As expected, hypoxia increased phosphorylation of an established AMPK target, acetyl-CoA carboxylase (ACC) in WT but not AMPK DKO cells, confirming AMPK activation in WT cells and loss of AMPK activity in DKO cells (Fig. 1C, D; fig. S1F, G).

Together, these results establish hypoxia as a physiological trigger for AMPK-dependent TFE3 nuclear translocation, mechanistically distinct from nutrient sensing.

### AMPK is required for hypoxia-induced lysosomal and mitochondrial biogenesis mediating cell survival

TFEB/3 drive expression of genes mediating autophagy, lysosomal function, and mitochondrial biogenesis, processes critical for surviving cellular stress (Malik *et al*., 2023; Martina *et al*., 2014; Martina and Puertollano, 2018; Yan *et al*., 2016). We have previously shown that FLCN regulates AMPK-dependent autophagy and stress survival in nematodes and mammalian cells (El-Houjeiri *et al*., 2019; Possik *et al*., 2015; Possik *et al*., 2014; Yan *et al*., 2014), and that FLCN loss induces mitochondrial biogenesis through PGC-1α/ERRα activation (Yan *et al*., 2016). We therefore assessed whether AMPK is required for these adaptive responses to hypoxia. Using the probe DQ-BSA, which fluoresces upon lysosomal degradation, we found increased lysosomal activity in WT MEFs after 2 hours of hypoxia, while AMPK DKO cells showed no increase (Fig. 1E, F). This indicates that AMPK is essential for hypoxia-induced lysosomal function.

We next evaluated mitochondrial biogenesis using flow cytometry to measure mitochondrial mass and membrane potential after 24 hours of hypoxia. WT cells showed significant increases in both mitochondrial mass and potential, whereas AMPK DKO cells exhibited no such changes (Fig. 1G-J). This is consistent with the requirement for AMPK and TFE3-mediated activation of PGC-1α/ERRα that has been previously described (Malik *et al*., 2023; Yan *et al*., 2016). Collectively, these results demonstrate that AMPK drives a coordinated program of lysosomal activation and mitochondrial expansion in response to low oxygen.

Given that lysosomal activity and healthy mitochondria are critical for stress survival, as we demonstrated in metabolic stress conditions in whole animals (Possik *et al*., 2015; Possik *et al*., 2014; Yan *et al*., 2014), we tested whether AMPK protects cells from hypoxia-induced cell death. After 12 hours of hypoxia, 85% of WT MEFs remained viable compared to only 65% of AMPK DKO MEFs; by 48 hours, viability decreased to 42% in WT versus 23% in AMPK DKO cells (Fig. 1K). Consistent with increased apoptosis, pro-Caspase 6 levels increased earlier in AMPK DKO cells, and cleaved Caspase-6 appeared at 12 hours in AMPK-deficient cells versus 24 hours in WT cells (fig. S2A-C). These findings establish that AMPK-mediated TFE3 activation is not merely correlated with but functionally required for survival under hypoxia.

### FLCN is required downstream of AMPK for hypoxic TFE3 regulation

The enhanced cytoplasmic retention of TFE3 in AMPK DKO cells during hypoxia (Fig. 1A, B) suggests that AMPK suppresses an activator of the mTORC1-TFEB/3 pathway. Since FLCN acts as a GAP for RagC/D to activate Rag heterodimers and mTORC1 (Lawrence et al., 2019; Shen et al., 2019; Tsun *et al*., 2013), and FLCN interacts with AMPK through FNIP1/2 (Baba *et al*., 2006; Hasumi *et al*., 2008; Takagi *et al*., 2008), we hypothesized that AMPK inhibits FLCN to release TFE3 from mTORC1-dependent suppression.

To test this, we generated AMPK/FLCN triple knockout (TKO) MEFs. Consistent with our prior work (Paquette *et al*., 2021), FLCN KO MEFs exhibited constitutive nuclear TFE3 regardless of oxygen levels (Fig. 2A, B). These findings are also consistent with chronic TFE3 activation when FLCN is absent (Hong et al., 2010). Critically, AMPK/FLCN TKO cells phenocopied FLCN KO cells: TFE3 remained nuclear under both normoxia and hypoxia (Fig. 2A, B). FLCN deletion did not impair AMPK activation during hypoxia (Fig. 2C) thus, confirming that FLCN acts downstream of AMPK. Furthermore, FLCN KO cells showed enhanced survival in hypoxia compared to WT cells (57% FLCN KO MEFs vs 42% WT MEFs) (Fig. 2D). These findings suggest that constitutive TFE3 activation provides a survival advantage. These results are consistent with the pro-survival phenotype conferred by FLCN loss in *C. elegans* under anoxia and other mitochondrial stressors which we previously reported (Possik *et al*., 2014). These genetic experiments demonstrate that FLCN is necessary for AMPK-dependent regulation of TFE3 during hypoxia.

**Fig. 2.**
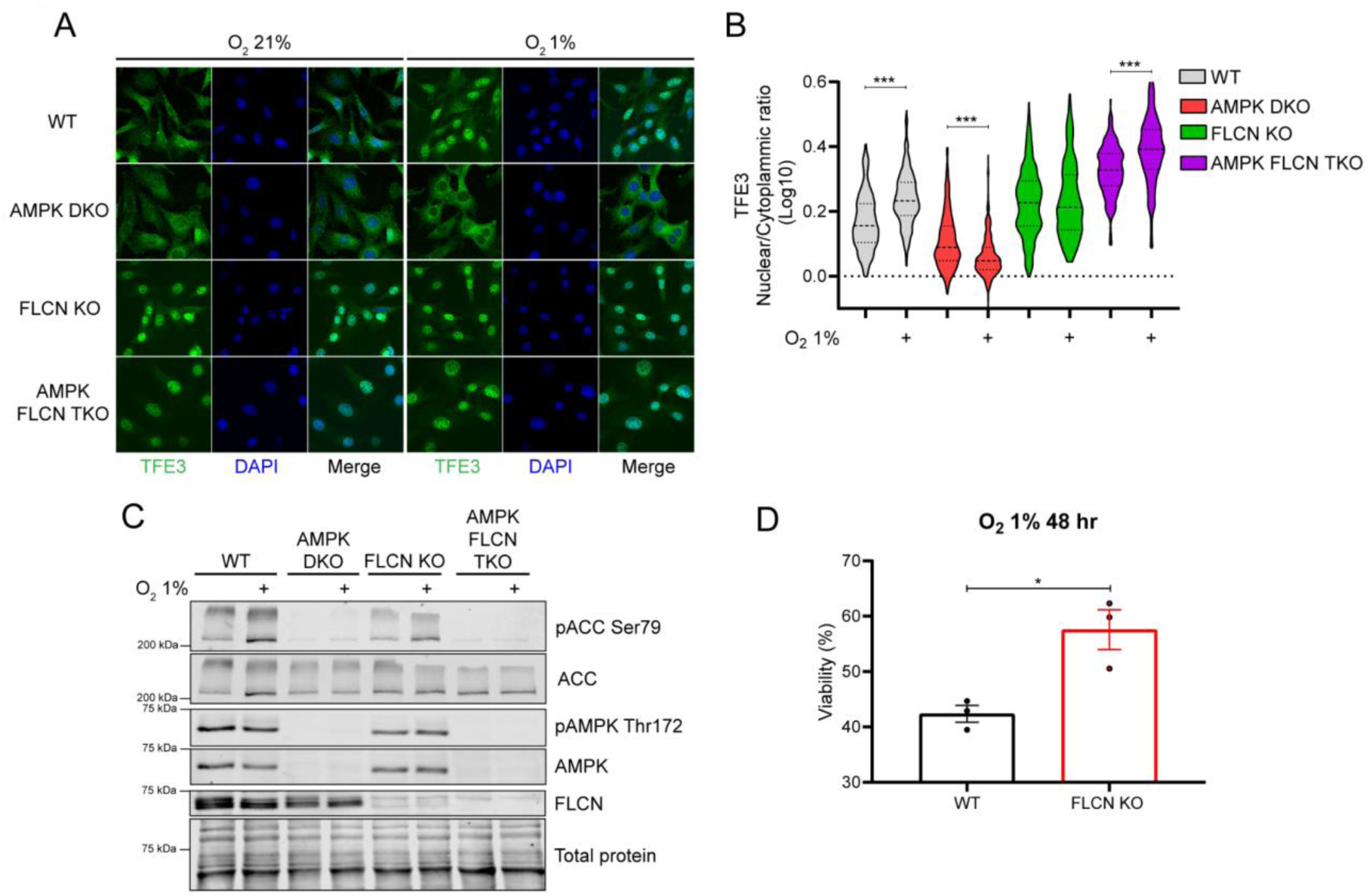
FLCN is downstream of AMPK for TFE3 regulation in hypoxia. **(A)** Representative immunofluorescence images of TFE3 localization in the indicated cell lines and conditions. (B) Nuclear/cytoplasmic ratio quantification of A. (C) Immunoblotting of AMPK activation in normoxic or hypoxic conditions in WT, AMPK DKO, FLCN KO and AMPK + FLCN TKO MEFs. (D) Cell viability in WT and FLCN KO MEFs incubated in hypoxia for 48 hours. Data are represented as the mean ± SEM. Three independent experiments were performed, at least 80 cells per immunofluorescence experiment and condition were examined, *p-value < 0.05, *** p-value < 0.001.

### AMPK phosphorylates FLCN at serine 130, inhibiting its GAP activity

To determine how AMPK regulates FLCN, we purified FLCN-FNIP1 complexes from AMPK-proficient and -deficient cells under normoxia or hypoxia followed by mass spectrometry. We identified FLCN serine 130 (S130) phosphorylation (Fig. 3A) as hypoxia-induced and AMPK-dependent: S130 was phosphorylated in WT cells under hypoxia but not normoxia, and this phosphorylation was reduced in AMPK DKO cells (table S1). S130 is highly conserved from humans to yeast (Fig. 3B, fig. S3A), suggesting the potential functional importance of this phosphoacceptor site.

**Fig. 3.**
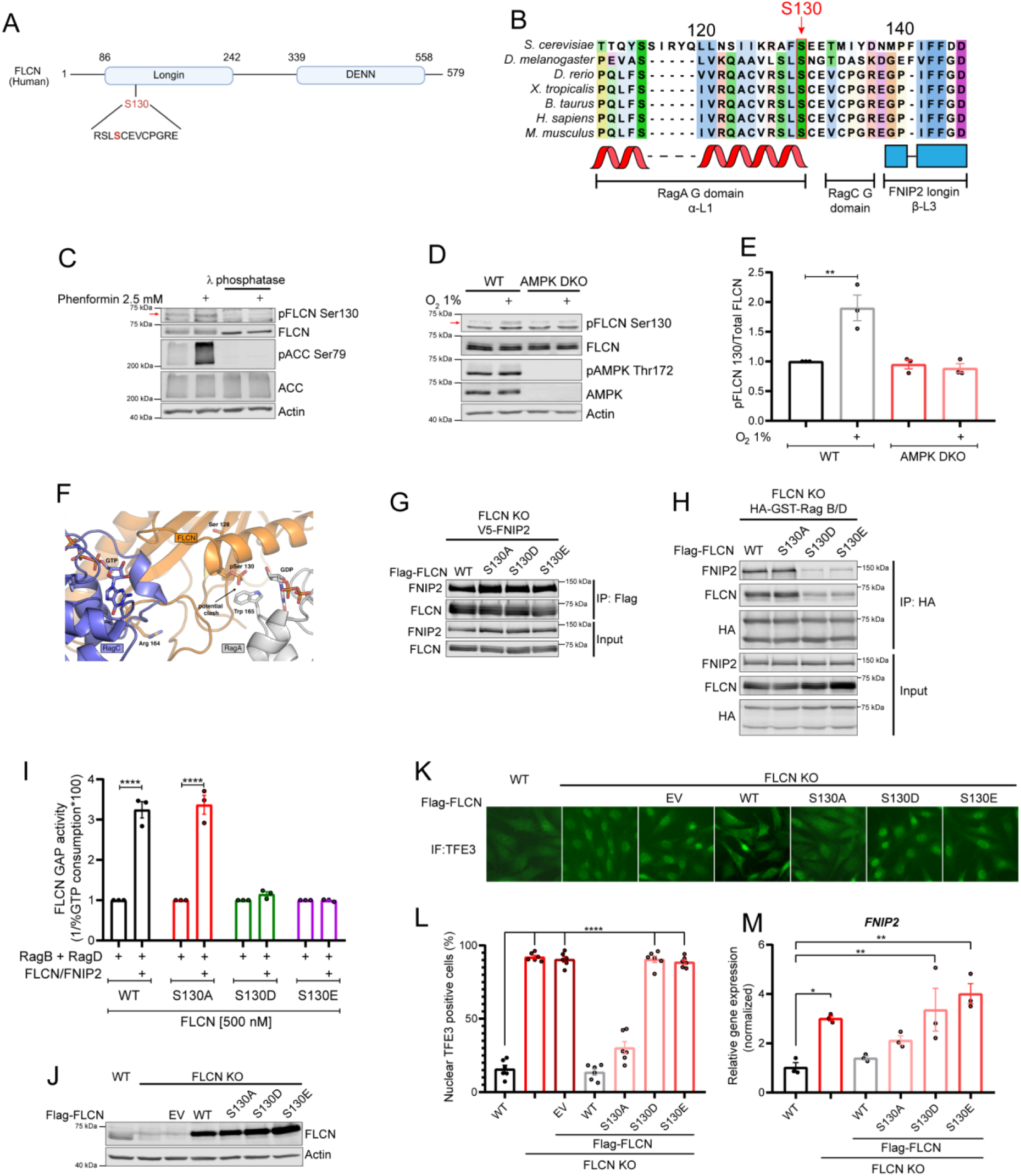
FLCN S130 phosphorylation inhibits GAP activity. **(A)** Graphical model of phosphorylated Ser130 in FLCN under hypoxia found by phosphoproteomics. (B) Sequence alignment of FLCN across species. (C) Representative immunoblotting of λ phosphatase incubation in cell extracts following phenformin treatment. (D) Representative immunoblotting of pFLCN Ser130 in WT or AMPK DKO cells incubated in normoxia or hypoxia. (E) Quantification of pFLCN Ser130 on Total FLCN ratio from D. (F) Model of Serine 130 phosphorylation function in the binding of FLCN with the Rag-GTPases. (G) Representative immunoblotting of Serine 130 WT, A, D or E Flag-FLCN mutants immunoprecipitation and their binding to FNIP2. (H) Representative immunoblotting of HA-GST-Rag B/D WT immunoprecipitation in the presence of Flag-FLCN constructs. (I) GAP activity of Flag-FLCN proteins in combination with GST-FNIP2 towards Rag-GTPases. (J) Representative immunoblotting of Flag-FLCN constructs stably expressed in FLCN KO MEFs. (K) Representative immunofluorescence images of TFE3 in the indicated cell lines. (L) Percentage of TFE3-nuclear-positive cells of K. (M) *FNIP2* mRNA expression analyzed by qPCR in the indicated cell lines. Three independent experiments were performed, at least 80 cells per immunofluorescence experiment were examined, *p-value < 0.05, **p-value < 0.01, **** p-value < 0.0001. Red arrow indicates specific pFLCN S130 band.

We generated a phospho-specific antibody against FLCN pS130 and confirmed its specificity by λ-phosphatase treatment (Fig. 3C). Hypoxia and pharmacological AMPK activation both induced S130 phosphorylation in WT but not AMPK DKO cells (Fig. 3D, E; fig. S3B, C). While S130 shows a partial match to the AMPK consensus motif (lacking the hydrophobic residue at -5 position) (Schaffer et al., 2015), our genetic data unambiguously demonstrate AMPK-dependent regulation of this site.

Structural modeling revealed that S130 lies in an α-helix within the RagA GTPase-binding interface of FLCN (Lawrence *et al*., 2019). Phosphorylation at this position would be predicted to create a steric clash, preventing FLCN-Rag interaction (Fig. 3F). To test this, we generated phosphomimetic (S130D, S130E) and non-phosphorylatable (S130A) FLCN mutants. Immunoprecipitation experiments showed that all mutants bound to FNIP2 at the level comparable to the WT (Fig. 3G). This indicates that the mutations on S130 do not disrupt FLCN-FNIP2 complex assembly. However, S130D and S130E showed dramatically reduced binding to Rag-GTPases compared to WT or S130A FLCN (Fig. 3H), confirming that S130 phosphorylation specifically disrupts the FLCN-Rag interaction.

Since FLCN must bind Rag-GTPases to exert its GAP activity, we predicted that S130 phosphomimetic mutants should be catalytically inactive. Using an *in vitro* GTPase assay, we found that WT and S130A FLCN (in complex with FNIP2) exhibited robust GAP activity toward RagC/D, while S130D and S130E mutants completely lacked activity (Fig. 3I). This demonstrates that S130 phosphorylation is a potent inhibitory switch for FLCN function.

To assess the functional implications of this phosphorylation, we rescued FLCN KO MEFs with WT, S130A, S130D, or S130E FLCN (Fig. 3J). Cells expressing S130D and S130E FLCN phosphomimetic mutants showed predominantly nuclear TFE3 and elevated expression of the TFEB/3 target gene FNIP2, phenocopying FLCN KO cells (Fig. 3K-M). In contrast, WT and S130A FLCN restored cytoplasmic TFE3 localization and normalized FNIP2 expression (Fig. 3K-M).

Finally, we identified multiple FNIP1 phosphorylation sites using mass spectrometry (including the five reported by Malik et al. plus an additional S594), but functional assays showed this phosphorylation event did not affect FLCN GAP activity (fig. S4A-C). This suggests that FNIP1 may have FLCN and GAP-independent functions (Xiao et al., 2024) that are regulated via these phosphorylation sites. Together, these results establish that FLCN S130 phosphorylation inhibits its GAP activity by preventing Rag-GTPase binding, thereby uncoupling TFE3 from mTORC1 control.

### The hypoxia response operates independently of GATOR1

GATOR1 is the GAP complex for RagA/B and is essential for mTORC1 inhibition during amino acid starvation (Bar-Peled *et al*., 2013). Building on our finding that the FLCN-AMPK-TFEB/3 axis can function independently of mTORC1 in the innate immune response in nematodes (El-Houjeiri *et al*., 2019), we investigated whether GATOR1 mediates the hypoxia response. We generated NPRL2 KO MEFs lacking the catalytic GATOR1 subunit. As expected, amino acid starvation failed to induce TFE3 nuclear translocation in NPRL2 KO cells (Fig. 4A-C), confirming the established role of GATOR1 in amino acid sensing. Remarkably, however, hypoxia induced robust TFE3 nuclear translocation in NPRL2 KO cells that was comparable to WT NPRL2 counterparts (Fig. 4A-C). This phenotype was confirmed in human cells (fig. S5A-D).

**Fig. 4.**
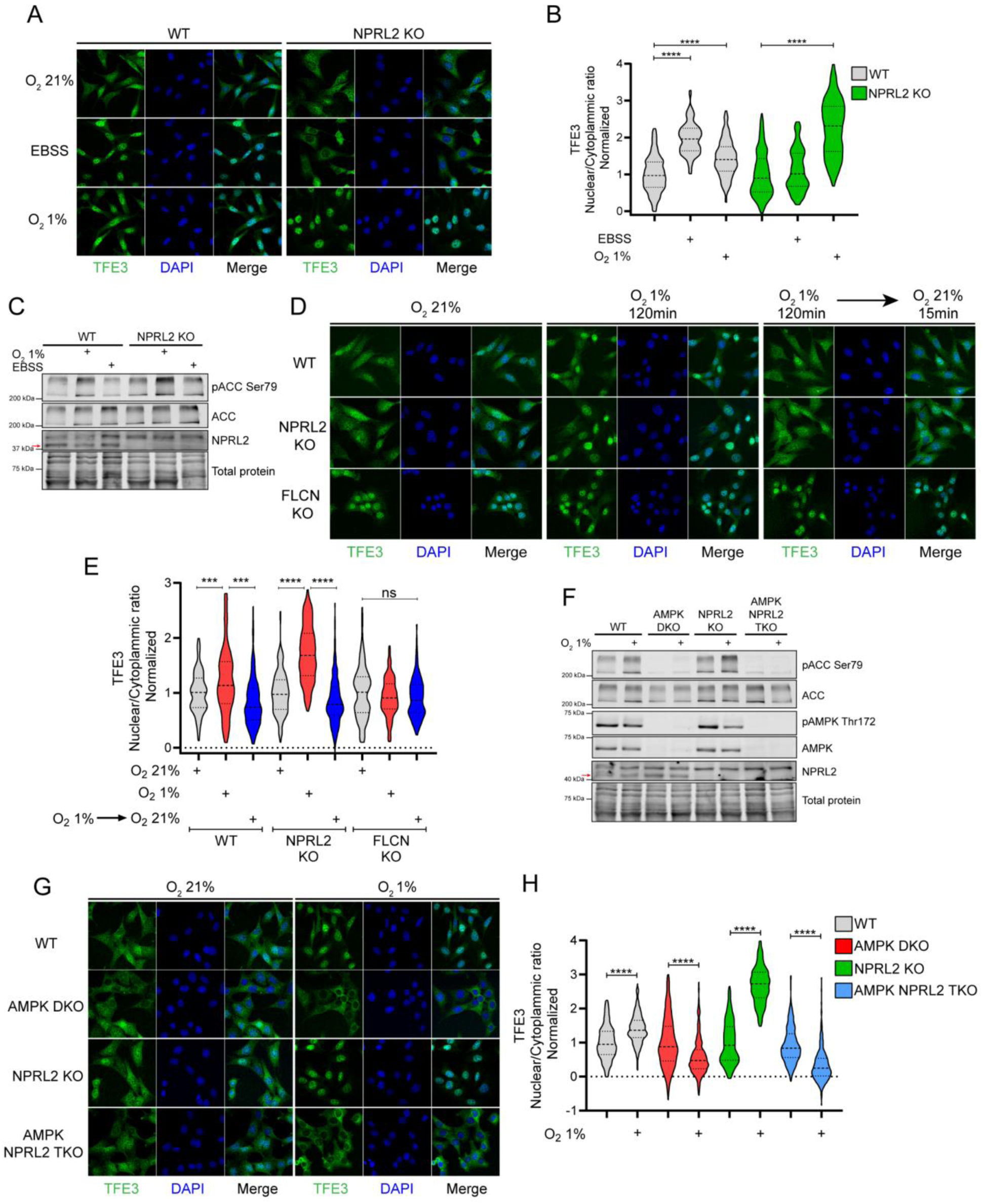
Hypoxia activates TFE3 independently of GATOR1. **(A)** Representative immunofluorescence images of TFE3 localization in the indicated cell lines and conditions. (B) Nuclear/cytoplasmic ratio quantification of A. Data are normalized to the control condition of each cell line. (C) Immunoblotting of AMPK activation upon two hours of hypoxia or one hour of starvation incubation of WT and NPRL2 KO MEFs. (D) Representative immunofluorescence images of TFE3 localization in the indicated cell lines in normal, hypoxia, or re-oxygenation conditions. (E) Nuclear/cytoplasmic ratio quantification of D. (F) Immunoblotting of AMPK activation in normoxic or hypoxic conditions in WT, AMPK DKO, NPRL2 KO and AMPK + NPRL2 TKO MEFs. (G) Representative immunofluorescence images of TFE3 localization in the indicated cell lines and conditions. (H) Nuclear/cytoplasmic ratio quantification of G. Data are normalized to the control condition of each cell line. Data are represented as the mean ± SEM. Three independent experiments were performed, at least 80 cells per immunofluorescence experiment and conditions were examined, *** p-value < 0.001, **** p-value < 0.0001, ns=not significant. The red arrow indicates the specific NPRL2 band.

We next examined whether TFE3 nuclear localization was dynamically regulated by oxygen availability. Cells were exposed to hypoxia for 120 minutes, then returned to normoxia for 15 minutes. TFE3 rapidly re-localized to the cytoplasm upon re-oxygenation in both WT and NPRL2 KO cells, but not in FLCN KO cells (Fig. 4D, E), demonstrating that FLCN is necessary for TFE3 cytoplasmic localization in an oxygen-dependent manner.

To test whether AMPK acts through this GATOR1-independent mechanism, we generated AMPK/NPRL2 triple knockout MEFs (Fig. 4F). In these cells, TFE3 remained cytoplasmic during hypoxia (Fig. 4G, H), phenocopying AMPK DKO cells rather than NPRL2 KO cells. This genetic epistasis experiment demonstrates that AMPK operates upstream of a GATOR1-independent pathway to inactivate Rag-GTPases during hypoxia.

Consistent with Rag inactivation, expression of constitutively active Rag mutants (RagB Q99L/RagD L77L) prevented TFE3 nuclear translocation during hypoxia (fig. S6A, B), demonstrating that Rag inactivation is necessary for the hypoxia response. These results establish oxygen-dependent Rag regulation as a mechanistically distinct signaling branch that acts in parallel to GATOR1 (fig. S6C).

### Hypoxia induces an AMPK-dependent TFE3 transcriptional program

Having established the mechanism of TFE3 activation by hypoxia, we performed RNA-seq in WT and AMPK DKO MEFs under normoxia and hypoxia to define the transcriptional consequences. Unbiased clustering identified seven gene clusters (fig. S7); two hypoxia-induced clusters were particularly informative. Cluster 6 comprised canonical hypoxia-response genes (HIF-1α targets involved in glycolysis and angiogenesis) that were induced independently of AMPK (Fig. 5A, B, D), consistent with our prior observations that FLCN loss promotes HIF-1α activation (El-Houjeiri *et al*., 2021). In contrast, Cluster 7 contained lysosomal genes previously identified as TFEB/3 targets—including *Flcn, Fnip2, Rragc, Mcoln1, Atp6v1c1, Slc38a7,* and *Gadd45a*—which were induced by hypoxia only in WT cells (Fig. 5A, C, D). qPCR validation confirmed AMPK-dependent induction of these genes with lysosomal functions (Fig. 5E).

**Fig. 5.**
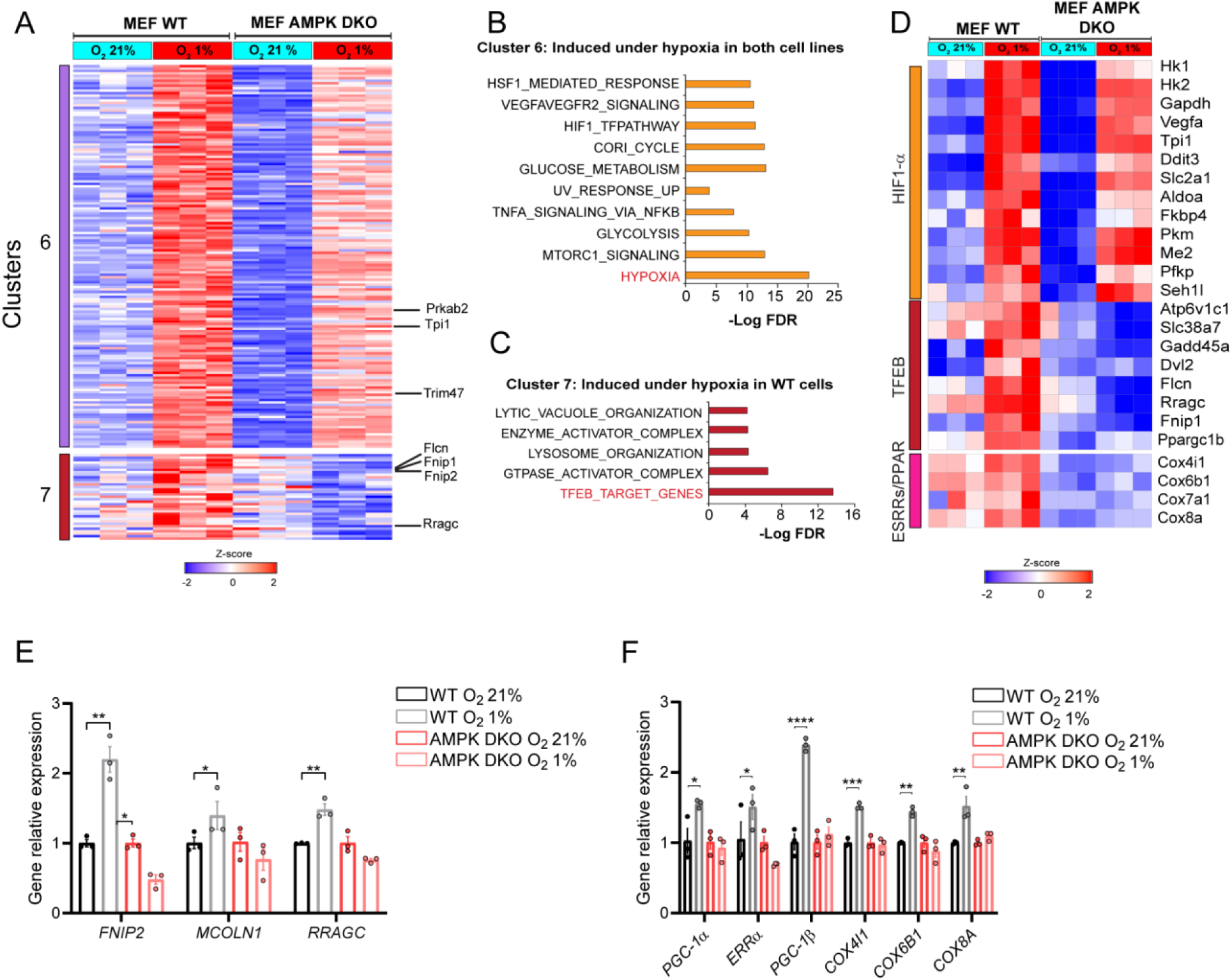
AMPK is required for hypoxia-induced TFEB/3 transcriptional program. **(A**) Unbiased hierarchical clustering of genes significantly modulated in response to hypoxia in one or both cell lines (FC ≥ 1.3, p value ≤ 0.05). (B and C) Transcription factors, pathways and hallmarks analysis of genes assigned to the indicated clusters using MSigDB. (D) Heatmap of AMPK-dependent and -independent regulated genes in response to hypoxia. (E and F) qPCR analysis of lysosomal-related genes (E) and mitochondrial-related genes (F). WT and AMPK DKO MEFs were incubated in normoxic or hypoxic conditions for 120 minutes prior to all experiments. Data are represented as the mean ± SEM. Three independent experiments were performed, *p-value < 0.05, **p-value < 0.01, *** p-value < 0.001, **** p-value < 0.0001.

Additionally, we identified AMPK-dependent induction of mitochondrial genes including *Ppargc1a* and *Ppargc1b* (encoding PGC-1α and PGC-1β), their transcriptional partner *Esrra* (ERRα), and downstream respiratory chain components (*Cox4i1, Cox6b1, Cox8a*) (Fig. 5D, F). This is consistent with our previous demonstration that FLCN loss leads to chronic AMPK activation and robust induction of PGC-1α/ERRα, driving mitochondrial biogenesis and oxidative metabolism (Yan *et al*., 2016). PGC-1α and β are established TFEB/3 targets that drive mitochondrial biogenesis (Betschinger et al., 2013; Malik *et al*., 2023; Wada et al., 2016), explaining the AMPK-dependent mitochondrial expansion we observed (Fig. 1G-J). These transcriptional data confirm that AMPK orchestrates a coordinated TFEB/3-driven program of lysosomal activation and mitochondrial biogenesis specifically in response to hypoxia, distinct from the HIF-1α-mediated glycolytic response.

### FLCN phosphorylation and TFE3 activation occur in hypoxic tumor regions and correlate with poor patient outcomes

Hypoxia is a hallmark of solid tumors, and both HIF-1α and TFEB/3 have been implicated in cancer progression (Bertout et al., 2008; Choudhry and Harris, 2018; Di Malta et al., 2023; Zhao et al., 2024). Building on our *in vitro* findings, showing TFE3 nuclear localization and transcriptional activation upon low oxygen tension, we examined whether TFE3 is activated in hypoxic tumor regions *in vivo*. We performed multiplex immunofluorescence for TFE3 and CA9 (a canonical hypoxia marker) from human breast cancer patient-derived xenografts (PDXs) (Savage et al., 2020). Strikingly, nuclear TFE3 was enriched in CA9-positive hypoxic areas, while it was predominantly cytoplasmic in CA9-negative regions (Fig. 6A-C). Next, we evaluated the transcriptional activity of TFEB/3 and HIF-1α in the RNA-seq data set of our breast cancer PDXs biobank (Savage *et al*., 2020) using curated TFEB/3 and HIF-1α signatures. We found a positive correlation between TFEB/3 and HIF-1α transcriptional activity (Fig. 6D). These findings support our *in vitro* findings, demonstrating TFE3 activation in the hypoxic tumor microenvironment.

**Fig. 6.**
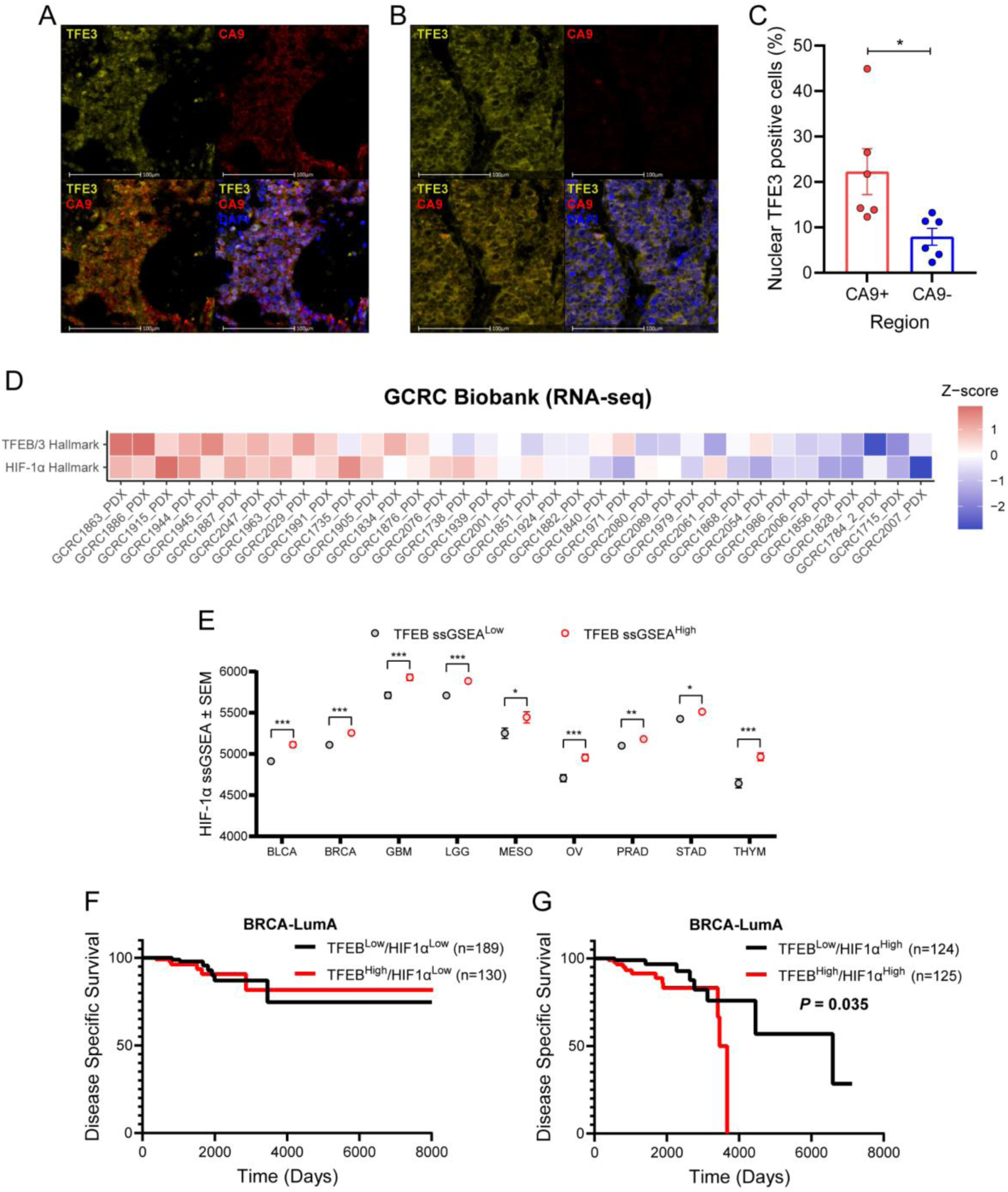
Hypoxia correlates with TFE3 activity in human tumors. (A and. **B)** Representative IHC images demonstrating TFE3 nuclear localization in hypoxic (A) or non-hypoxic (B) regions of breast cancer PDXs. (C) Quantification of the percentage of cells with nuclear TFE3 in CA9-positive or negative regions. (D) mRNA heatmap of the TFEB/3 and HIF-1α hallmarks in 36 breast cancer PDXs. (E) TFEB/3 and HIF-1α gene signature enrichment across several cancers from the PAN CAN TCGA dataset. (F and G) Kaplan-Meier curves showing disease-specific survival among Luminal A breast cancer patients with high or low TFEB/3 activity and either low (F) or high (G) HIF-1α activity. Data are represented as the mean ± SEM, *p-value < 0.05, **p-value < 0.01, ***p-value < 0.001.

To assess the clinical relevance of TFEB/3 and HIF-1α co-activation in cancer patients, we performed single-sample gene set enrichment analysis (ssGSEA) on the TCGA PanCancer dataset. The two gene signatures positively correlated across nine cancer types: bladder cancer, breast cancer, thymoma, prostate adenocarcinoma, glioma (both lower-grade and glioblastoma), ovarian cancer, mesothelioma, and gastric adenocarcinoma (Fig. 6E). Most importantly, luminal A breast cancer patients with high expression of TFEB/3 signature showed significantly reduced disease-specific survival specifically when HIF-1α signature is elevated as well (Fig. 6F, G), demonstrating that both elevated signatures remodel a relatively easy-to-treat cancer type into a highly aggressive cancer.

These observations suggest that tumors activate the hypoxia-AMPK-FLCN-TFE3 pathway to survive in low-oxygen environments. While HIF-1α promotes angiogenesis and metabolic reprogramming (Rankin et al., 2016; Zhao *et al*., 2024), TFE3 activation may provide complementary survival signals through lysosomal activity and mitochondrial expansion and quality control. The association with poor outcomes identifies this pathway as a potential therapeutic target in hypoxic tumors.

## Discussion

The mTORC1-TFEB/3 axis has been mainly studied in the context of amino acid availability, where GATOR1 inactivates the Rag-GTPases upon nutrient starvation, leading to mTORC1 dissociation from lysosomes and TFEB/3 dephosphorylation (Martina and Puertollano, 2013; Settembre *et al*., 2012). Our work reveals that oxygen limitation activates the Rag-GTPases-mTORC1-TFEB/3 branch through an entirely different mechanism: FLCN inhibition by AMPK phosphorylation, which prevents Rag activation regardless of amino acid status. This emerges as an oxygen-specific pathway since other AMPK activators, such as glucose starvation, induce TFE3 nuclear translocation via other means (Li et al., 2018).

The independence from GATOR1 is particularly significant and suggests that cells have evolved parallel inputs into the Rag system: GATOR1 for amino acid sensing and FLCN inhibition for oxygen detection.

We propose that the existence of independent branches lies in the differential cellular needs imposed by distinct stresses and stress response plasticity (Chen et al., 2025). For instance, amino acid starvation primarily requires proteolytic capacity to recycle proteins for their building blocks, hence autophagy and lysosomal activation are paramount. Hypoxia, while also requiring autophagy, additionally demands a mitochondrial quality control: damaged mitochondria (due to ROS accumulation) must be removed through mitophagy, while new and functional mitochondria must be generated and expanded to ensure hypoxic adaptation and cell survival (Gutsaeva et al., 2008; Kung-Chun Chiu et al., 2019; Tohme et al., 2017).

Our finding of FLCN S130 as a regulatory phosphorylation site addresses a major gap left by recent work (Malik *et al*., 2023). It was demonstrated that pharmacological AMPK activation induces FNIP1 phosphorylation at five serines, which was proposed to regulate FLCN function, though no direct evidence for altered FLCN GAP activity was shown (Malik *et al*., 2023).

Moreover, this study relied on pharmacological AMPK activators, thereby obscuring the physiological relevance of AMPK-mediated FLCN regulation and leaving it unaddressed. We now show that (i) hypoxia is a physiological activator of the AMPK/TFEB/3 axis; (ii) FLCN, not just FNIP1, is phosphorylated in response to hypoxic AMPK activation; (iii) FLCN S130 phosphorylation directly inhibits GAP activity by preventing Rag binding; and (iv) this mechanism is functionally required for survival under hypoxia. Nevertheless, although we found that FNIP1 phosphorylation has no role in FLCN GAP activity regulation, FNIP1’s function independently of the FLCN-TFEB/3 axis should be further studied (Xiao *et al*., 2024).

Located in the Rag-binding interface, the S130 site is particularly well-suited to act as a regulatory switch (Lawrence *et al*., 2019). Its phosphorylation creates a steric hindrance that prevents FLCN from engaging its substrate (RagC/D). This model suggests a rapid and reversible inactivation: upon oxygen restoration, S130 dephosphorylation would enable more FLCN to bind Rags and reactivate mTORC1, as we observed TFE3 nuclear exclusion in re-oxygenation experiments. Additionally, the evolutionary conservation of S130 from yeast to human suggests that this regulatory mechanism is ancient and fundamental for FLCN function.

While our genetic data established that S130 phosphorylation is regulated by AMPK, the molecular mechanisms remain to be investigated. The S130 sequence shows a partial match to the AMPK consensus motif (lacking a hydrophobic residue at the -5 position) (Schaffer *et al*., 2015). However, whether AMPK directly phosphorylates this site or acts through an intermediate kinase remains an open question.

AMPK activation under hypoxia is mechanistically interesting. The classic model relies on ATP depletion increasing the AMP/ATP ratio, which allosterically activates AMPK, and promotes its phosphorylation by upstream kinases as well (Gowans et al., 2013; Suter et al., 2006). However, we found that hypoxia decreased ATP, ADP, and AMP levels without changing their ratios These findings are consistent with previous reports (Mungai *et al*., 2011). It was demonstrated that hypoxia activates AMPK primarily through mitochondrial ROS production and calcium release rather than through changes in adenine nucleotides (Emerling et al., 2009; Hinchy et al., 2018; Mungai *et al*., 2011). This suggests that AMPK senses the production of mitochondrial ROS during hypoxia, distinct from the energy depletion commonly observed under nutrient withdrawal (Gwinn *et al*., 2008). This mechanistic distinction further underscores the specificity of different stress pathways converging on AMPK.

Our current findings integrate with and complement our extensive prior reports on the FLCN-AMPK axis. We previously showed that FLCN regulates autophagy and is required for metabolic stress survival in an AMPK-dependent manner in animals and cells (Possik *et al*., 2015; Possik *et al*., 2014), and that FLCN loss leads to chronic AMPK activation, driving PGC-1α/ERRα-mediated mitochondrial biogenesis and oxidative metabolism (Yan *et al*., 2016; Yan *et al*., 2014). These effects have critical repercussions in tumor progression. We have demonstrated that FLCN loss promotes metabolic rewiring, including activation of HIF-1α-dependent glycolysis and angiogenesis (El-Houjeiri *et al*., 2021). Results of our current work reveal the mechanistic basis for these observations: hypoxia-induced AMPK phosphorylation of FLCN S130 creates a regulatory switch that coordinates autophagy, lysosomal function, and mitochondrial biogenesis through TFEB/3 activation. This mechanism is likely to be implicated in both physiological adaptation to hypoxia as well as pathologies wherein oxygen sensing plays a prominent role, including cancer.

Our finding that hypoxia induces PGC-1α/β expression in an AMPK-dependent manner directly parallels our earlier demonstration that FLCN loss (which mimics FLCN phosphorylation) induces these same transcriptional coactivators (Yan *et al*., 2016). This confirms that the AMPK→FLCN→TFE3→PGC-1α pathway we have defined represents a central regulatory axis controlling mitochondrial adaptation to stress. The importance of this pathway is underscored by our observation that AMPK-deficient cells fail to mount both lysosomal activity and mitochondrial biogenesis responses to hypoxia, resulting in significantly impaired survival.

Solid tumors grow in environments characterized by both nutrient limitation and hypoxia (Ciepla and Smolarczyk, 2024). Our discovery that these stresses activate TFEB/3 through distinct mechanisms helps explain tumor adaptation. Our previous work showed that FLCN, FNIP1, and FNIP2 are downregulated in many human cancers, particularly aggressive basal-like breast carcinomas, and that FLCN loss in luminal breast cancer promotes tumor growth through TFE3-dependent induction of the Warburg effect, oxidative phosphorylation, angiogenesis and lysosomal biogenesis (El-Houjeiri *et al*., 2021). The current findings suggest that even when FLCN protein is present, tumors can functionally inactivate it through AMPK-mediated phosphorylation, gaining the same proliferative advantages without requiring genetic mutations.

Critically, our discovery of FLCN inhibition proposes a molecular mechanism for how tumors might exploit the hypoxia response: rather than requiring loss-of-function mutations in FLCN (as in BHD syndrome) (Schmidt and Linehan, 2018), tumors might functionally inactivate FLCN through phosphorylation, gaining the survival advantages of constitutive TFEB/3 activation while retaining other FLCN functions. This mechanism may also explain the tissue specificity of BHD syndrome. Germline FLCN mutations predispose to renal tumors, particularly in the chronically hypoxic renal medulla where oxygen levels are normally 1-2% (Brezis and Rosen, 1995; Liu *et al*., 2022). Importantly, renal tumors in BHD patients are mainly hybrid oncocytic tumors and chromophobe renal cell carcinomas (Schmidt and Linehan, 2018). These tumor types are characterized by an accumulation of mitochondrial content (Klomp et al., 2010). Our model predicts that in this environment, WT FLCN would be constitutively phosphorylated and inactive. Cells with germline FLCN loss lack functional enzyme activity even under normoxic conditions, conferring a selective advantage during hypoxic episodes while allowing damage to accumulate and thereby enhancing their tumorigenic potential.

While HIF-1α drives glycolysis and angiogenesis to address hypoxia (Zhao *et al*., 2024), the AMPK-FLCN-TFE3 pathway provides complementary survival signals: autophagy to generate nutrients from damaged proteins and organelles, and mitochondrial biogenesis to restore respiratory capacity when oxygen becomes transiently available in fluctuating tumor microenvironments (Lai et al., 2016). The correlation between TFEB/3 and HIF-1α signatures across multiple cancer types, and their association with poor patient outcomes, suggests that co-activation of these pathways marks particularly aggressive hypoxic tumors.

From a therapeutic perspective, targeting the AMPK-FLCN-TFE3 axis in hypoxic tumors presents both opportunities and challenges. mTOR inhibitors and AMPK activators have shown limited efficacy in most solid tumors (Jeon and Hay, 2015; Roczniak-Ferguson et al., 2012; Settembre *et al*., 2012), possibly because they inhibit all mTORC1 substrates indiscriminately. In long term, these findings may open new therapeutic avenues by selectively blocking the TFEB/3 branch—perhaps by preventing FLCN phosphorylation or inhibiting TFEB/3 activity (Lin et al., 2023)—might prove more effective in hypoxic, HIF-1α-high tumors.

Finally, *in vivo* validation using recently developed TFEB/3 activity reporter mice (Brunialti et al., 2024) will be essential to understand this pathway’s physiological roles during exercise (which creates transient muscle hypoxia), high-altitude exposure, and tumor development. Such studies may reveal whether pharmacological modulation of this pathway has therapeutic potential for ischemic diseases or cancer.

## Conclusion

Taken together, we have identified an oxygen-dependent regulation of TFEB/3. Mechanistically, it is a distinct signaling branch separated from the well-established nutrient-sensing pathway. By phosphorylating FLCN at S130, AMPK creates a molecular switch that inactivates FLCN’s GAP activity, releasing TFE3 from mTORC1 control to activate a transcriptional program essential for hypoxia survival. This pathway, which builds upon our extensive characterization of FLCN-AMPK-TFEB/3 signaling (El-Houjeiri *et al*., 2021; El-Houjeiri *et al*., 2019; Paquette *et al*., 2021; Possik *et al*., 2015; Possik *et al*., 2014; Yan *et al*., 2016; Yan *et al*., 2014), is present in tumors with hypoxic microenvironments and correlates with poor patient outcomes, suggesting new therapeutic opportunities. Overall, our findings demonstrate how cells specifically respond to different signals through modular organization: distinct entry points (amino acid sensors-GATOR1 vs. AMPK-FLCN) into a common signaling system (Rag-GTPases). It allows appropriate responses in context-dependent scenarios. This principle likely extends to other signaling networks where specificity emerges from regulated access points rather than separate pathways.

## Acknowledgments

We thank Dr. Benoit Viollet for AMPK DKO MEFs, Dr. Laura Schmidt for Flcn-floxed mice. The flow cytometry work/ cell sorting was performed in the Flow Cytometry Core Facility for flow cytometry and single-cell analysis of the Life Science Complex and supported by funding from the Canadian Foundation for Innovation. Images were collected and/or image processing and analysis for this manuscript was performed in the McGill University Advanced BioImaging Facility (ABIF), RRID:SCR_017697. Data analyses were enabled by compute and storage resources provided by the Digital Research Alliance of Canada.

## Funding

This work was supported by grants to AP:

The Kidney Foundation of Canada, Terry Fox Foundation (TFF-166128). CIHR (PJT-165829).

The Cancer Research Society.

JM.JRR was supported by CONAHCYT (SECIHTI) (CVU 818450).

## Author contributions

Conceptualization: JMJRR and AP

Methodology: JMJRR, APB, SD, SH and CLK.

Investigation: JMJRR, APB, SD, SH, CLK, JUS, MP, BN and AP

Visualization: JMJRR, SD, JUS and AP

Supervision: AP

Writing – original draft: JMJRR and AP

All authors reviewed and approved the final manuscript

## Competing interests

The authors declare no competing interests.

## Data, code, and materials availability

RNA-seq data have been deposited in GEO under accession GSE325779. All other data are available in the main text or supplementary materials. Reagents are available from the corresponding author upon reasonable request and completion of a material transfer agreement.

## Supplementary Materials

Figs. S1 to S7

Tables S1 to S2

## Materials and Methods

### Cell lines cultures and treatments

Primary MEFs were isolated from C57BL/6 E12.5 Flcn floxed mice (generously provided by Dr. L.S. Schmidt, NCI, Bethesda, MD, USA) or Flcn/Prkaa1/Prkaa2 floxed mice and cultured as described previously. FLCN and FLCN/AMPK KO MEFs were generated as described previously (Paquette *et al*., 2021). WT and AMPK DKO MEFs cells were generously provided by Dr. Benoit Viollet (Institut Cochin INSERM, Paris, France). FLCN KO MEFs were transduced with retrovirus with Flag-FLCN WT or S130 mutants, and Vector only with Flag was used as control. Transduced cells were selected with Blasticidin S HCl (2 ug/mL, Gibco, cat#A11139-03). AMPK DKO HEK293T cells were generated and tested previously (Paquette *et al*., 2021). HEK293T stably overexpressing CA Flag-RagB+Flag-RagD were transduced with lentivirus and selected with puromycin. Cell lines were maintained in Dulbecco’s modified Eagle’s medium (DMEM) (Wisent, cat# 319-005CL) supplemented with 10% fetal bovine serum (FBS) (Wisent, cat# 080-150), 100 U/ml penicillin+100 µg/ml streptomycin (Wisent, cat# 450-201-EL), and 50 µg/mL gentamycin (Wisent, cat# 450-135) in 5% CO_2_ at 37°C. For EBSS starvation experiments, cells were washed twice with PBS and incubated in Earle’s Balanced Salt Solution (EBSS, Sigma, cat# E2888) for the indicated time. For drug treatment experiments, cells were incubated for 1 hour in medium containing one of the following reagents: Dimethyl sulfoxide (DMSO) (0.1%, Bioshop Canada, cat# DMS666), AICAR (2 mM, Enzo Life Sciences, cat# 89158-090), Phenformin (2.5 mM, Sigma, cat#P7045), 991 (50 µM, Glixx Laboratories Inc, cat# GLXC-09267) and 2-Deoxy-D-Glucose (10 mM, Sigma, cat#D6134). The final DMSO concentration never exceeded 0.1% and this concentration was shown to have no detrimental effect on all the studied cells. For hypoxia experiments, cells were incubated in 1% O_2_, 5% CO_2_ at 37°C for the indicated times.

### CRISPR-generated KO cells

NPRL2 KO and AMPK+NPRL2 TKO MEFs were generated by transduction with LentiCRISPR-Cas9-derived virus containing the mouse-NPRL2 gRNA sequence 5’-CAGCGAGTTCCACCCAACGC-3’. NPRL2 KO HEK293T cells were generated by transiently transfecting cells with LentiCRISPER-Cas9 with the human-NPRL2 gRNA sequence 5’-GTGATCTTGGGTCCCAGCGT-3’. FLCN KO HEK293T cells were generated using the reported human-FLCN sgRNA sequence 5’-TCGCACATGTCCGACTTTTT-3’. Cells were selected with puromycin (2 mg/ml) and single cell cloned. Control cells were infected or transfected with non-coding sgRNA LentiCRISPR-Cas9.

### Plasmids

FNIP1 WT or 5SA or 5SE constructs were codon-optimized and cloned in pCAG-GST-Flag or pCW57-Flag vectors. Codon-optimized pCAG-TWIN-STREP-Flag-FLCN (Addgene #164981) and pCAG-GST-FNIP2 (Addgene #164982) were previously reported (Lawrence *et al*., 2019). V5-FNIP2 was cloned in pcDNA3 vector. Flag-FLCN was cloned in pCW57 Tet off vector. FLCN Ser130 mutants were generated by directed mutagenesis. Other plasmids include: pRK5-HA-GST-RagB Wt (Addgene #19301), pRK5-HA-GST-RagD Wt (Addgene #19307), pLJM1-Flag-RagB wt (Addgene #19313), pLJM1-Flag-RagB 99L (Addgene #19315), pLJM1-Flag-RagD wt (Addgene #19316), pLJM1-Flag-RagD 77L (Addgene #19317).

### Transient transfection

HEK293T cells at 75% confluence were transfected with Polyethylenimine 5 μg/μg of DNA (Polysciences, cat# 23966). For FNIP1-FLCN purification and Mass spectrometry, 8 μg of each codon-optimized vectors were transfected. For FLCN-FNIP2 immunoprecipitation experiments, 1 μg of V5-FNIP2 and 100 ng of Flag-FLCN constructs were used. For Rag-GTPases-FLCN/FNIP2 IP experiments, 150 ng of HA-GST-RagB and 150 ng of HA-GST-RagD were transfected with V5-FNIP2 and Flag-FLCN. For FLCN GAP activity, 4 ug of GST-FNIP1 or GST-FNIP2 plus 8 ug of Flag-FLCN were transfected and 8 ug HA-GST-RagB and 8 ug HA-GST-RagD were transfected for Rags enrichment. For FNIP1-FLCN immunoprecipitation and Rag-FLCN/FNIP1 immunoprecipitation, 250 ng GST-FNIP1 and the previously mentioned DNA amounts were transfected. All experiments were performed 24 hours post-transfection.

### Antibodies

TFE3 (clone F3X8T; Cell Signaling Technology, cat# 19950S) AMPKα (Cell Signaling Technology, cat# 2532), p-AMPKα (Thr172) (Cell Signaling Technology, cat# 2531), Actin (Sigma, Cat# MAB1501), HA (Covance, cat# MMS-101R), FLCN (Cell Signaling Technology, cat# 3697), FNIP2 (Sigma, cat# HPA042779), FNIP1 (Abcam, cat# ab134969), ACC (Cell Signaling Technology, cat# 3676), phospho-ACC (Ser79) (Cell Signaling Technology, cat# 3661), Caspase 6 (Cell Signaling Technology, cat# 9762), Cleaved Caspase 6 Asp162 (Cell Signaling Technology, cat# 9761), Flag M2 ( Sigma, cat# F3165), Lamin A (Santa Cruz, cat# sc-71481), Tubulin ( Sigma, cat# T9026), Revert™ 700 Total Protein Stain for Western Blot Normalization (LI-COR, cat# 926-11016), antibodies are commercially available. Anti-mouse-FLCN was generated by the McGill animal resource center services. Anti-phospho-FLCN Ser 130 was developed by Capralogics Inc. (MA, USA).

### Protein extraction and immunoblotting

Cells were washed twice with PBS, lysed in RIPA-light lysis buffer (50mM Tris-HCl pH 8.0, 150mM NaCl, 5mM EDTA, 1% NP40, 0.1% SDS, 0,1% Triton X-100, 5 mM NaF, 0.1 mM Sodium orthovanadate (Na_3_VO_4_), 1 mM benzamidine, 5 mM Sodium pyrophosphate (NaPPi), supplemented with a complete protease inhibitor cocktail (Roche, cat# 5056489001), and cell lysates were cleared by centrifugation at 13000 x g. Proteins were separated on SDS-PAGE gels and revealed by western blot using the antibodies listed above in the Odyssey Classic Imaging System (LI-COR).

### Lambda-phosphatase assay

Cells were washed twice with PBS, lysed in Triton lysis buffer (20mM Tris pH 7.5, 150mM NaCl, 1% Triton X-100 (Bioshop, cat# TRX506), supplemented with a complete protease inhibitor cocktail (Roche, cat# 5056489001), and cell lysates were cleared by centrifugation at 13000 x g. 50 mg of protein was incubated with 400 Lambda protein phosphatase units in lysis buffer supplemented with 1mM MnCl_2_. Samples were incubated at 30 °C for 2 hours. 2X Laemmli sample buffer was added to the samples and boiled for 5 min at 95 °C. Proteins were separated on SDS-PAGE gels and revealed by western blot using the antibodies listed above.

### Immunofluorescence

Cells were gently washed with PBS and fixed in petri dishes with 3.7% formaldehyde (Bioshop Canada, cat# FOR201.500). After fixation, cells were washed twice with PBS and then permeabilized with 0.3% Triton X-100 (Bioshop Canada, cat# TRX506.100). Cells were incubated with TFE3 primary antibody in 5% BSA in PBS for O/N at 4°C. Cells were washed three times with PBS and incubated with the corresponding secondary antibodies conjugated to Alexa Fluor 488 (Thermo Fisher, cat# A-11008), Alexa Fluor ® 555 Phalloidin (88 nM, Cell Signaling Technology, cat# 8953) and DAPI (2 μg/ml) (Molecular Probes, cat# D-1306) in 5% BSA in PBS for two hours at 22°C. PBS-washed cover slips were mounted and observed with LSM 800 Confocal Microscope (Zeiss). Nuclear-Cytoplasmic ratio score was performed using CellProfiler software (https://cellprofiler.org/) (Stirling et al., 2021), DAPI signal was used to generate a Nuclear mask and Phalloidin signal was used to design a Cytoplasm mask. TFE3-DAPI Pearson’s correlation was evaluated using Fiji (Schindelin et al., 2012).

### Multiplex fluorescent immunohistochemistry

Staining on formalin-fixed, paraffin-embedded tumor sections derived PDXs was performed following the manufacturer’s instructions for Opal™ Multiplex Immunofluorescence (Akoya Biosciences, Opal Assay Development Guide), with minor modifications. Briefly, sections were deparaffinized and rehydrated using successive 3-minute incubations in histological-grade xylene (3X; Sigma, 534056-4L) and decreasing concentrations of ethanol (100% ethanol, 3X 3 min; 95% ethanol, 1X 3 min; 70% ethanol, 1X 3 min), followed by 3–5 washes in distilled water.

Antigen retrieval was then performed in citrate buffer (pH 6.0, 10× Antigen Retriever; Millipore Sigma, C9999-1000ML) using a pressure cooker for 10 minutes. Sections were incubated in 3% hydrogen peroxide for 10 minutes and blocked using eBioscience™ IHC/ICC High Protein Blocking Buffer (Thermo Fisher Scientific, 00-4952-54) for 5 minutes. Slides were rinsed once with 2% BSA-TBST.

Anti-TFE3 antibody was diluted in TBST containing 2% BSA (1:100) and incubated for 30 minutes at room temperature. Slides were washed 3X in TBST and incubated with ImmPRESS HRP-conjugated anti-rabbit polymer secondary antibody (Vector Laboratories, VECTMP740150) for 30 minutes at room temperature. Slides were washed 3X in TBST. Fluorescent signal was developed using Opal 520 tyramide signal amplification reagent (Akoya Biosciences, OP-001001) for 10 minutes at room temperature.

After a second round of antigen retrieval and blocking, anti-CA9 antibody (Sigma-Aldrich, 379R-16) was diluted in TBST containing 2% BSA (1:100) and incubated for 1 hour at room temperature. Sections were washed 3X in TBST and incubated with ImmPRESS HRP-conjugated anti-mouse polymer secondary antibody (Vector Laboratories, VECTMP745250) for 30 minutes at room temperature. Slides were washed 3X in TBST. Fluorescent signal was developed using Opal 570 reagent (Akoya Biosciences, OP-001003) for 10 minutes. At the end of the experiment, sections were counterstained with DAPI and mounted for fluorescence imaging. Quantification of staining was performed using Halo image analysis software (Indica Labs).

### Co-immunoprecipitation

For Rags immunoprecipitation, transfected HEK293T cells were lysed in CHAPS lysis buffer (40mM Hepes pH 7.9, 0.3% CHAPS (Bioshop, cat# CHA003), 2.5mM MgCl_2_) supplemented with a complete protease inhibitor cocktail (Roche, cat# 5056489001), 4 mM NaF, 1 mM Na_3_VO_4_, 0.2 mg/ml PMSF, 2.5 mM NaPPi, and put on ice for 20 min after one wash with PBS. Cell lysates were cleared by centrifugation at 13,000 x g for 10 min at 4°C and the lysate pre-cleared for 1 h with protein A/G magnetic beads (Thermo Scientific, cat# 88802). The cell lysates were immunoprecipitated overnight at 4°C using HA magnetic beads (Pierce, cat# PI88837). Beads were washed three times with lysis buffer supplemented with 150 mM NaCl and PMSF. Immunoprecipitations were eluted twice (and combined) in 2X Laemmli sample buffer and separated on SDS-PAGE gels and revealed by western blot using the antibodies listed above.

For FLCN-FNIP1 or FLCN-FNIP2 experiments, transfected HEK293T cells were washed with PBS once and lysed in Triton lysis buffer (20mM Tris pH 7.5, 150 mM NaCl_2_, 0.7% Triton X-100, 10% Glycerol, supplemented with a complete protease inhibitor cocktail (Roche, cat# 5056489001), 4 mM NaF, 2.5 mM NaPPi, cell lysates were incubated on ice for 20 minutes. Cell lysates were cleared by centrifugation at 13,000 x g for 10 min at 4°C. The cell lysates were immunoprecipitated overnight at 4°C using Flag M2 magnetic beads (Sigma, cat# M8823). Immunoprecipitations were eluted twice (and combined) in 2X Laemmli sample buffer and separated on SDS-PAGE gels and revealed by western blot using the antibodies listed above.

### GST pull-down and elution

HEK293T cells transfected with the indicated vectors were lysed with Triton lysis buffer after one PBS wash. Cell lysates were incubated at 4°C with gentle shaking for 20 minutes and cleared by centrifugation at 13,000 x g for 10 min at 4°C. Lysates were incubated with pre-washed glutathione-sepharose 4 FastFlow beads (GE Healthcare, 17075601) overnight at 4°C. Beads were washed with ice-cold PBS and incubated with 50 mM Tris-HCl, pH 8.0, 10 mM glutathione (Bioshop Canada, GTH001.5) for 30 minutes at 4°C with gentle shaking. Eluted proteins were cleared by centrifugation.

### *In vitro* GAP activity assay

15 % of eluted GST-FNIP1/ 2 + Flag-FLCN or HA-GST-RagB/D was loaded in an SDS-PAGE 8%. Coomassie staining was performed for Flag-FLCN and HA-GST-RagB/D quantification using a BSA-based curve. Different Flag-FLCN concentrations or a maximum of 500 mM were used, and a total of 500 mM of Rags were used for all experiments. FLCN GAP activity was tested using GTPase-Glo Assay (Promega, cat# V7681) following supplier’s instructions. Luminescence was measured using Fluostar Omage (BMG Labtech) directly in the plates.

### RT-qPCR

Total mRNA was isolated and purified using PuroSPIN total RNA Purification Kit (Luna Nanotech, cat#NK051). 1 μg of RNA was used for the reverse transcription step using the iScript Reverse Transcription Supermix for RT-qPCR (BioRad, 1708841). qPCR experiments were performed using the SYBR Green qPCR supermix (BioRad, 1725125) and specific primers (table S2) in an AriaMX Real-time PCR system (Agilent Technologies, USA). Relative expression of mRNAs was determined after normalization against the housekeeping gene *Tbp*.

### RNA extraction, RNA-sequencing and bioinformatics analysis

For RNA-seq experiments, total RNA from WT or AMPK MEFs incubated as indicated was isolated using TRIzol (Invitrogen 15596026). Total RNA was quantified, and its integrity was assessed using 5K / RNA / Charge Variant Assay Lab Chip and RNA Assay Reagent Kit (Perkin Elmer). Libraries were generated from 100 ng of total RNA as following: mRNA enrichment was performed using the NEBNext Poly(A) Magnetic Isolation Module (New England BioLabs). cDNA synthesis was achieved with NEBNext UltraExpress RNA Library Prep Kit (New England BioLabs) as per the manufacturer’s recommendations. Adapters and PCR primers were purchased from New England BioLabs. Libraries were quantified using the KAPA Library Quantification Kits - Complete kit (Universal) (Kapa Biosystems). Average fragment size was determined using a Fragment Analyzer 5300 (Agilent) instrument. The libraries were normalized and pooled and then denatured in 0.02N NaOH and neutralized using pre-load buffer. The pool was loaded at 170pM on an Illumina NovaSeq X Plus 25B lane following the manufacturer’s recommendations. The run was performed for 2×100 cycles (paired-end mode). A phiX library was used as a control and mixed with libraries at 1% level. Program BCL Convert 4.2.4 was then used to demultiplex samples and generate fastq reads. An alignment index for mm10 was prepared with STAR (Dobin et al., 2013) through the RSEM tool. The corresponding transcript annotation (Ensembl 98) in gtf format was provided to allow for transcripts quantification.

STAR’s splice junction database overhang parameter was set to 99 nucleotides. The raw FASTQ reads were aligned to the mm10 genome with STAR through the RSEM tool. Reverse strandedness was specified, in accordance with the stranded protocol used to generate the RNA-Seq libraries. The analysis produced both a genome and a transcript alignment (in BAM format), as well as per sample gene and isoform level expression tables. Cohort-wide count tables were created by combining each sample’s independent expression files. Additional QCs were performed on the raw counts to identify potential outliers and sample biases. These analyses were performed in the R statistical software environment using the DESeq2 (Love et al., 2014) and pheatmap libraries. The Z-score heatmap was constructed using the R package ComplexHeatmap (Gu, 2022). The gene sets for each cluster were subsequently submitted to EnrichR (Kuleshov et al., 2016), which is freely available at http://amp.pharm.mssm.edu/Enrichr/ (Kuleshov *et al*., 2016). The following libraries were used for the analysis: GO_2025, Reactome_2024, and CHEA_2022 libraries. Significant gene signatures were determined using the FDR q-val <0.05 cut-off.

### RNA-seq - PAN CAN TCGA patients

TFEB/3 signature (*FLCN, RRAGC, RRAGD, GADD45A, SLC38A7, FNIP2, MCOLN1, ATP6V1C1, FNIP1, SNX8, SQSTM1, PPARGC1A, PPARGC1B, LAMP1, LAMP2, CTSD, CTSB, NPC1, BECN1, ATG7, ARL8B, LIPA, CPT1A, TFEB*) and HIF-1α signature (*EGLN3, SLC2A1, SLC2A3, SLC16A3, HK1, HK2, PGK1, PKM2, CA9, VEGFA, LDHA, ANGPTL4, ANGPT2, LOX, LOXL2, TWIST1, TFRC, ADM, DDIT4, ALDOA, BNIP3, SERPINE1*) were used for further analysis.

RNA-seq normalized gene counts from all patients of the 33 TCGA cancer types were used to compute the ssGSEA scores for the TFEB/3 and HIF-1α signatures. The gsva function from the GSVA v2.0.7 bioconductor package (Hanzelmann et al., 2013) with a ssgseaParam object and normalization set to FALSE was used to generate the ssGSEA scores for each patient. Low and high ssGSEA were defined as bottom and top 40% of the patients, respectively.

### Viability Sulforhodamine B (SRB) assay

Cells were plated at 5 x 10^3^ or 2 x 10^3^ cell density per well in a 96-well plate and incubated in normoxia or hypoxia for 12 hours or 48 hours. Viability was evaluated using the SRB Assay / Sulforhodamine B Assay Kit (Abcam, cat# ab235935) following the manufacturer’s instructions. Absorbance at 565 nm was measured using Fluostar Omage (BMG Labtech).

### Mitochondrial content analysis

Sub-confluent cells were incubated with MitoTracker Red CMXRos (200 nM, Invitrogen, cat# M46750) and MitoTracker Green FM (20 nM, Invitrogen cat# M7512) in FBS-free DMEM media for 30 minutes at 37°C after the indicated treatments. Cells were collected and resuspended in PBS supplemented with FBS 2%. Fluorescent intensity was acquired with a Fortessa BD Biosciences cytometer. Data were analyzed with Flowing Software 2.5.1 and Floreada Cytometry.

### Lysosomal activity

MEFs were incubated with 5 μg/mL DQ Red BSA (Thermo Fisher, cat# D12051) for 2 h and washed twice with PBS. Treatments were applied for the indicated time. Cells were then collected, fixed, washed and resuspended in PBS. Fluorescent intensity was acquired with a Fortessa BD Biosciences cytometer. Data were analyzed with Flowing Software 2.5.1 and Floreada Cytometry.

### Mass Spectrometry analysis

Immunoprecipitated Flag-FNIP1 and Flag-FLCN were electrophoretically separated in a Mini-PROTEAN TGX Gels (Bio-Rad, cat# 4561094). Excised gel bands were cut into approximately 1 mm^3^ pieces. The samples were reduced with 1 mM DTT for 30 minutes at 60°C and then alkylated with 5mM iodoacetamide for 15 minutes in the dark at room temperature. Gel pieces were then subjected to a modified in-gel trypsin digestion procedure (Shevchenko et al., 1996).

Gel pieces were washed and dehydrated with acetonitrile for 10 min. followed by removal of acetonitrile. Pieces were then completely dried in a speed-vac. Rehydration of the gel pieces was with 50 mM ammonium bicarbonate solution containing 12.5 ng/µl modified sequencing-grade trypsin (Promega, Madison, WI) at 4°C. Samples were then placed in a 37°C room overnight.

Peptides were later extracted by removing the ammonium bicarbonate solution, followed by one wash with a solution containing 50% acetonitrile and 1% formic acid. The extracts were then dried in a speed-vac (∼1 hr). The samples were then stored at 4°C until analysis.

On the day of analysis, the samples were reconstituted in 5 - 10 µl of HPLC solvent A (2.5% acetonitrile, 0.1% formic acid). A nano-scale reverse-phase HPLC capillary column was created by packing 2.6 µm C18 spherical silica beads into a fused silica capillary (100 µm inner diameter x ∼30 cm length) with a flame-drawn tip (Peng and Gygi, 2001). After equilibrating the column each sample was loaded via a Famos auto sampler (LC Packings, San Francisco CA) onto the column. A gradient was formed and peptides were eluted with increasing concentrations of solvent B (97.5% acetonitrile, 0.1% formic acid).

As each peptide was eluted, they were subjected to electrospray ionization and then they entered into an LTQ Orbitrap Velos Pro ion-trap mass spectrometer (Thermo Fisher Scientific, San Jose, CA). Eluting peptides were detected, isolated, and fragmented to produce a tandem mass spectrum of specific fragment ions for each peptide. Peptide sequences (and hence protein identity) were determined by matching protein or translated nucleotide databases with the acquired fragmentation pattern by the software program, Sequest (ThermoFinnigan, San Jose, CA) (Eng et al., 1994). The modification of 79.9663 mass units to serine, threonine, and tyrosine was included in the database searches to determine phosphopeptides. Phosphorylation assignments were determined by the Ascore algorithm (Beausoleil et al., 2006). All databases include a reversed version of all the sequences and the data was filtered to between a one and two percent peptide false discovery rate.

### Statistical analysis

Data are expressed as mean ±SEM. All experiments and measurements were performed at least three times as indicated. Statistical analyses for all data were performed using student’s t-test, one-way ANOVA or two-way ANOVA as indicated using GraphPad Prism 9 software. Statistical significance is indicated in figures (*P<0.05, **P<0.01, ***P<0.001, ****P<0.0001).

## Notes

### Competing Interest Statement

The authors have declared no competing interest.

