## Supplementary Materials for "AMPK-dependent FLCN phosphorylation activates a survival response to hypoxia"

Figure S1.

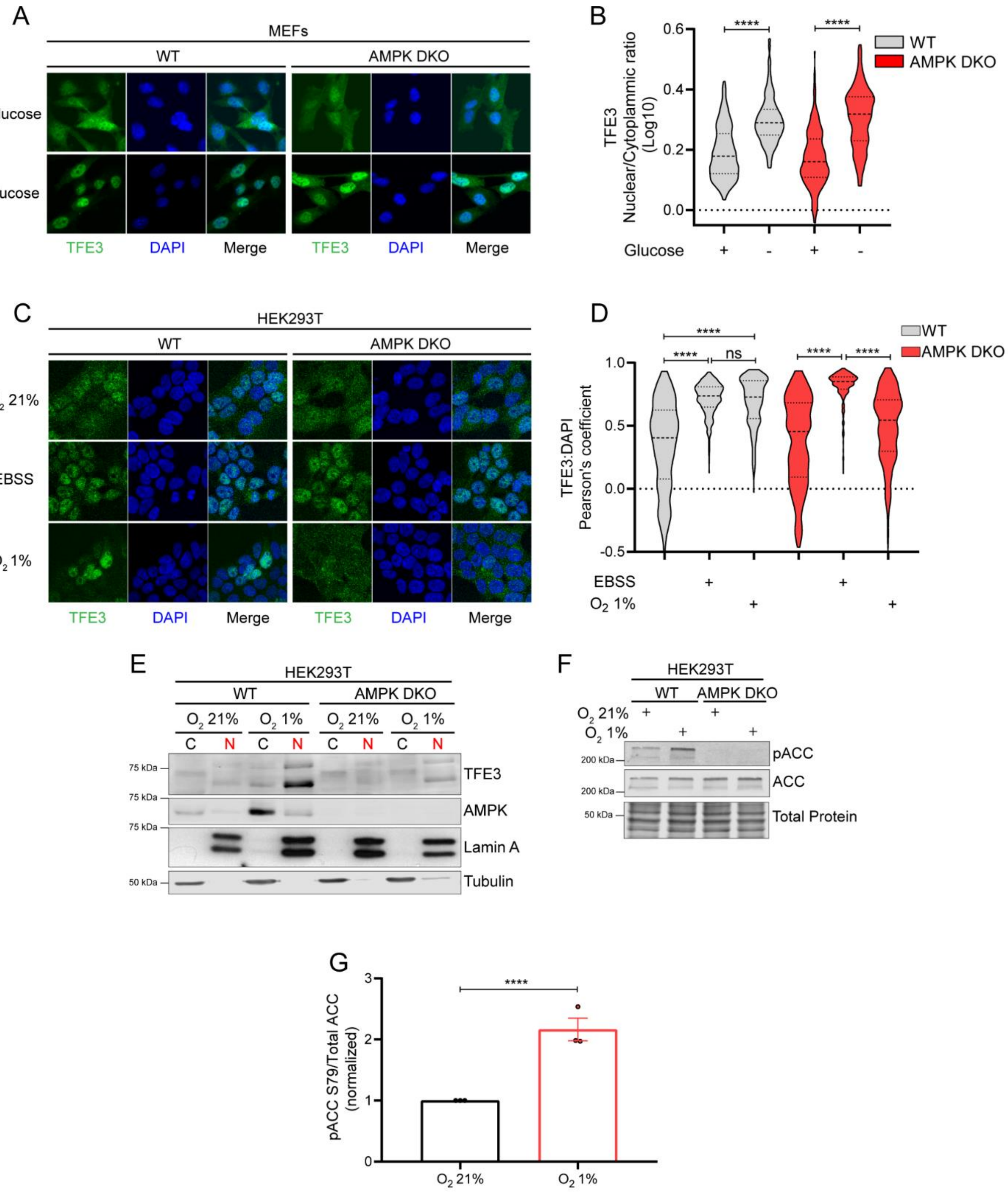


**Fig. S1. TFE3 translocation upon hypoxia depends on AMPK in human cells.**
**(A)** Representative immunofluorescence images of TFE3 localization in the indicated cell lines incubated in media with or without glucose. (B) Nuclear/cytoplasmic ratio quantification of A. (C) Representative immunofluorescence images of TFE3 localization in the indicated cell lines incubated in normal, starvation or hypoxia conditions. (D) Pearson's correlation of TFE3 signal

with DAPI signal of C. (E) Immunoblotting of cellular fractionation in WT and AMPK DKO HEK293T under normoxia or hypoxia. (F) Immunoblotting of AMPK target upon normal, starvation or hypoxic conditions in WT and AMPK DKO HEK293T cells. (G) Phospho-ratio quantification of AMPK target from D. Data are represented as the mean ± SEM. Three independent experiments were performed, at least 80 cells per immunofluorescence experiment and condition were examined, **** p-value < 0.0001, ns=not significant.

Figure S2.

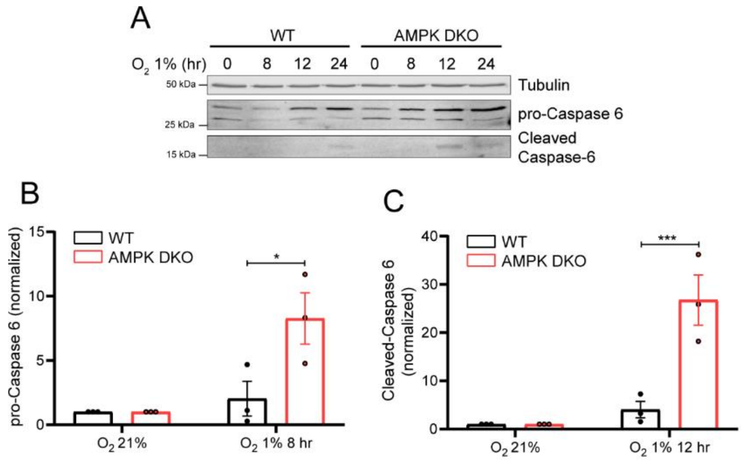


**Fig. S2. AMPK is needed to delay apoptosis in hypoxia.**
**(A)** Representative immunoblotting of pro-Caspase 6 and Cleaved Caspase 6 in the indicated cell lines incubated in hypoxia for the indicated times. (B and C) Quantification of pro-Caspase 6 (B) and Cleaved Caspase 6 (C) of the experiments from A. Data are represented as the mean ± SEM. Three independent experiments were performed, *p-value < 0.05, *** p-value < 0.001.

Figure S3.

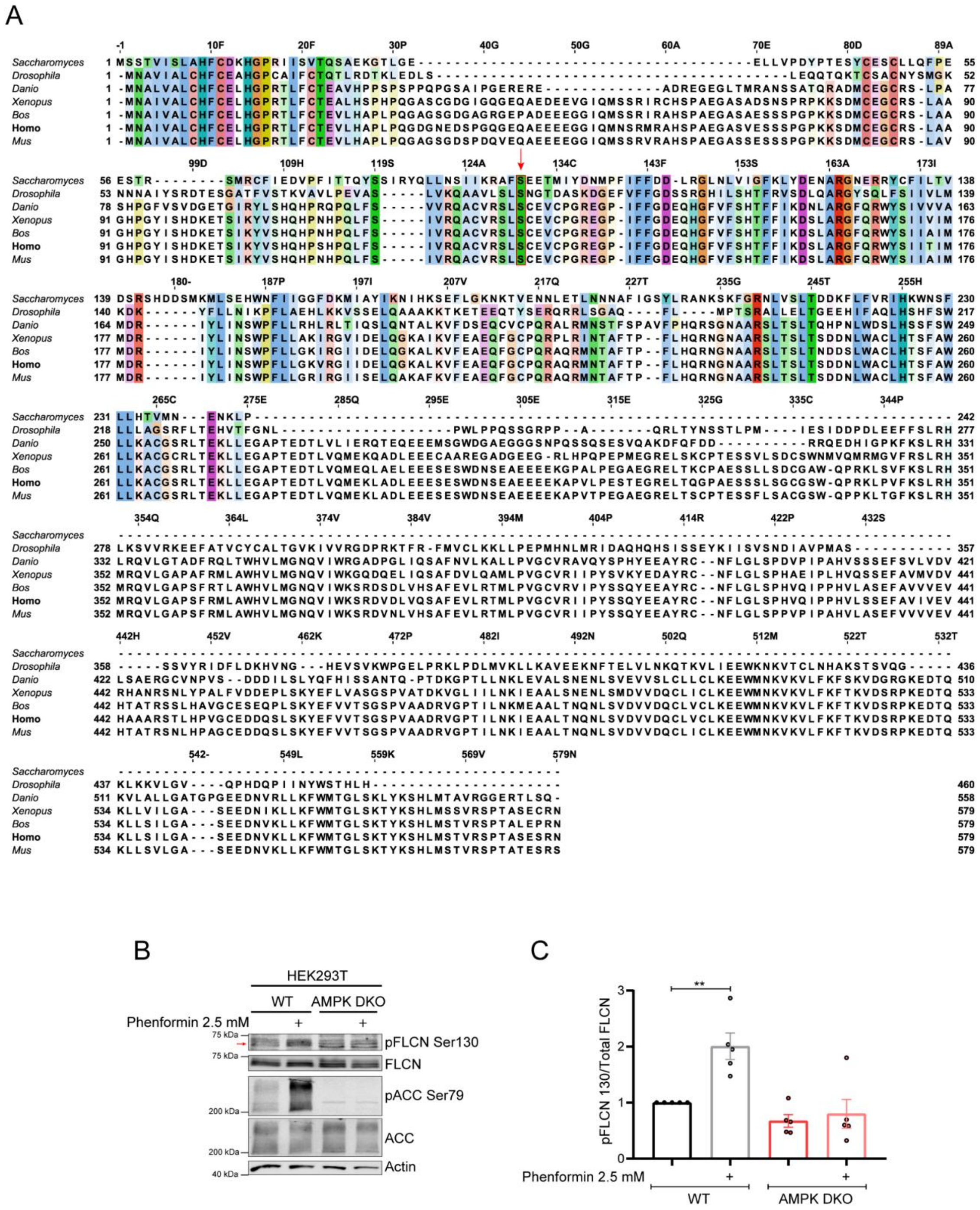


**Fig. S3. FLCN is phosphorylated upon AMPK activation.**

**(A)** Sequence alignment of FLCN of diverse species performed in T-COFFEE platform (https://tcoffee.crg.eu/), Ser130 is indicated by the red arrow and box. (B) Representative immunoblotting of pFLCN Ser130 and AMPK markers from cells treated with or without Phenformin 2.5 mM for 60 minutes. (C) Quantification of pFLCN Ser130 on Total FLCN ratio from C. Data are represented as the mean ± SEM. Three independent experiments were performed. ** p-value < 0.01. Red arrow indicates specific pFLCN S130 band.

Figure S4.

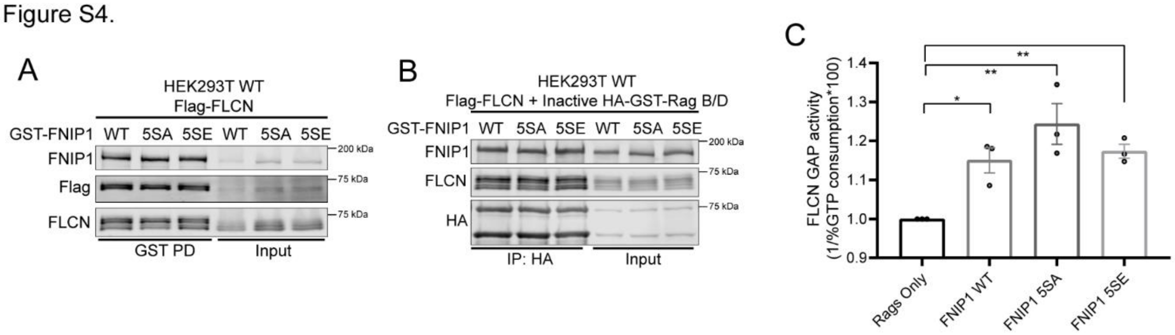


**Fig. S4. FNIP1 phosphorylation does not affect FLCN activity**

**(A)** Representative immunoblotting of GST-FNIP1 WT or 5SA or 5SE mutants pull-down and their binding to FLCN. (B) Representative immunoblotting of HA-GST-Rag B/D Inactive mutants immunoprecipitation in the presence of GST-FNIP1 constructs and Flag-FLCN. (C) GAP activity capacity of Flag-FLCN in combination with GST-FNIP1 constructs towards Rag-GTPases. Data are represented as the mean ± SEM. Three independent experiments were performed, * p-value < 0.05, ** p-value < 0.01.

Figure S5.

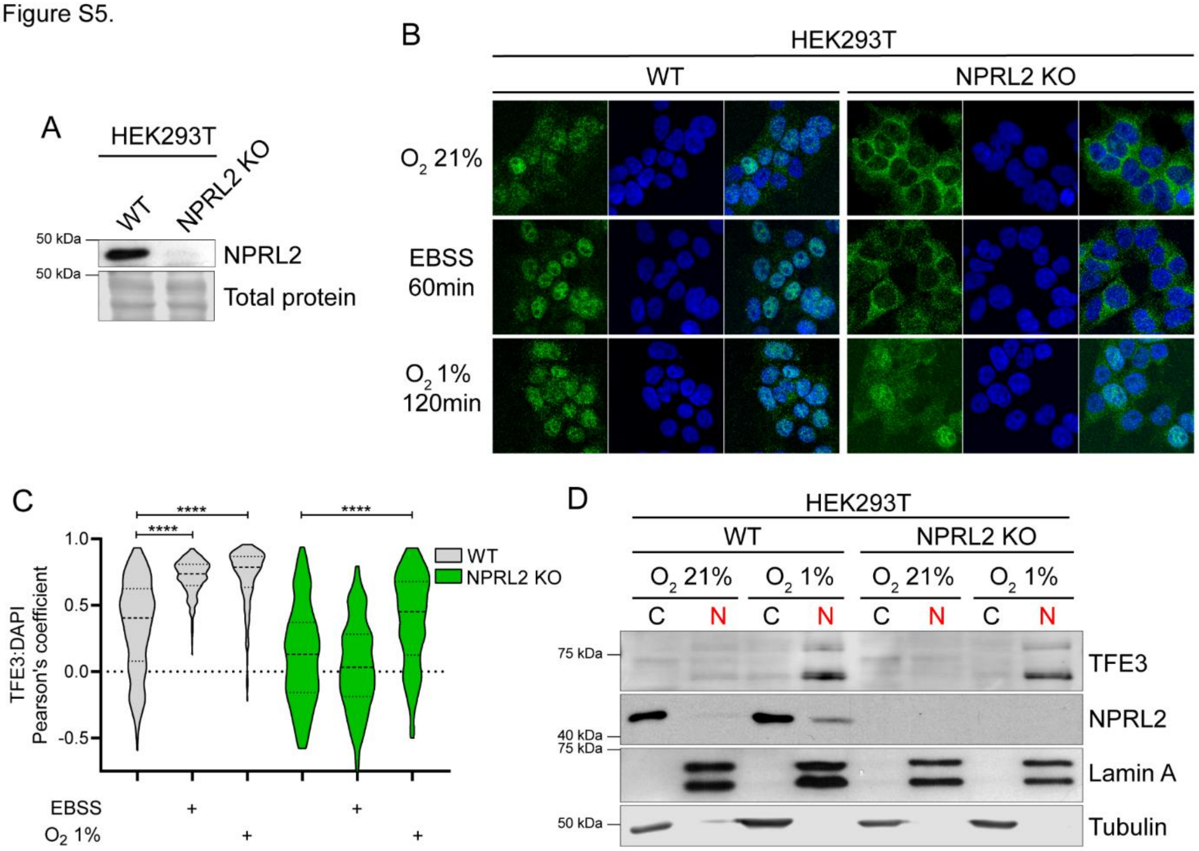


**Fig. S5. NPRL2 is dispensable for TFE3 translocation in response to AMPK activation and oxygen levels in human cells.**

**(A)** Representative immunoblotting of NPRL2 KO HEK293T cells generation. (B) Representative immunofluorescence images of TFE3 localization in the indicated cell lines incubated in normal, starvation or hypoxia conditions. (C) Pearson's correlation of B. (D) Immunoblotting of cellular fractionation in WT and NPRL2 KO HEK293T under normoxia or hypoxia. Data are represented as the mean ± SEM. Three independent experiments were performed, at least 80 cells per immunofluorescence experiment and condition were examined, **** p-value < 0.0001, ns=not significant.

Figure S6

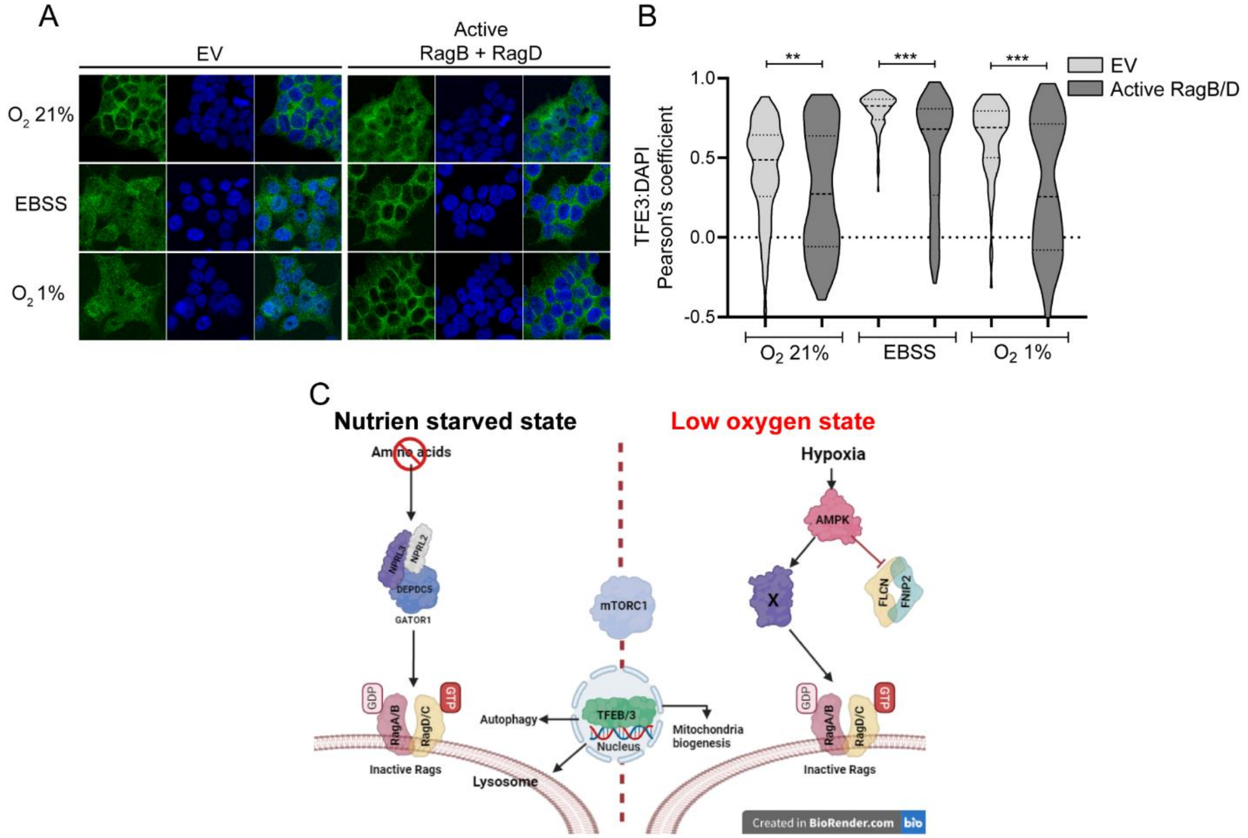


**Fig. S6. The nucleotide state of the Rag-GTPases controls TFE3 localization in response to hypoxia.**

**(A)** Representative immunofluorescence images of TFE3 localization in the indicated cell lines and conditions. (B) Pearson's correlation quantification of A. (C) Model of TFEB/3 localization and activity in response to amino acid starvation, which is AMPK independent, and under low-oxygen conditions where AMPK inhibits FLCN GAP activity, promoting the inactivation of the Rag-GTPases and TFEB/3 dephosphorylation and translocation to induce the expression of lysosomal and mitochondrial-related genes to contest cell death and encourage adaptation in response to hypoxia. Data are represented as the mean ± SEM. Three independent experiments were performed, at least 80 cells per immunofluorescence experiment and condition were examined, *p-value < 0.05, **p-value < 0.01, ***p-value < 0.001.

Figure S7.

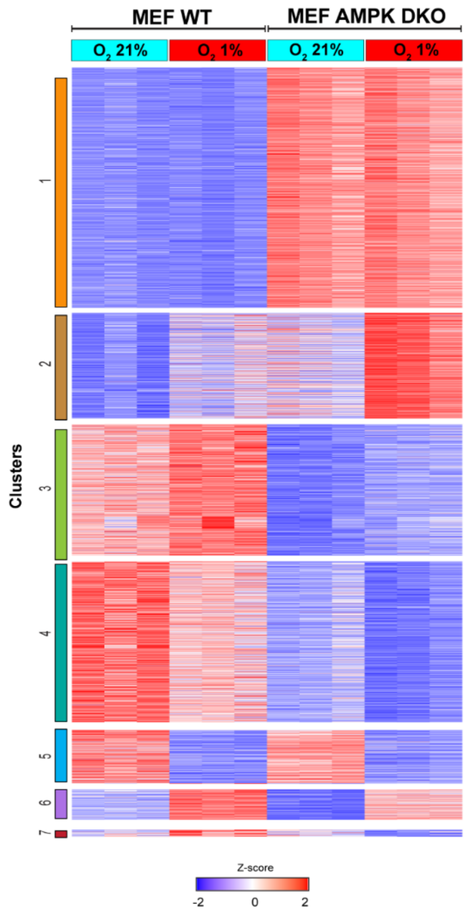


**Fig. S7. RNA-seq analysis from AMPK-sufficient and AMPK-deficient cells in response to hypoxia.**

**Table S1. (separate file)**
**FLCN and FNIP1 phosphopeptides.**

**Table S2. Primers for qPCR experiments.**

| **Gene** | **Forward** | **Reverse** |
|---|---|---|
| ***FNIP2*** | CCACACCTGGTGATAGGACTGTT | GCACCAAATGTCACAGAAAAATG |
| ***MCOLN1*** | CATGTTTGTGACGTTCGCGG | AGTACCTCCTGGGTGCTTGA |
| ***RRAGC*** | GCAACTTCACACACTCTGCG | CAGTGGGAAGGGTATCAGCC |
| ***COX4IL*** | CCTGATTCCCGCGATGCT | TCTCTTGCCAATCAGGCTCA |
| ***COX6B1*** | AACGCTACTCCGGGACAATC | AGTCTGGTTCTGGTTGGGGA |
| ***COX8A*** | TAGACCACTTTTGCCAGCCC | CCACCAAGCAGAGCCAATAC |
| ***PPARGC1A*** | CGGAAATCATATCCAACCAG | TGAGGACCGCTAGCA AGTTTG |
| ***PPARGC1B*** | ACTGAAAGAGGCCCAGCAGA | ATTGGAAGGGCCTTGTCTGA |
| ***ESRRA*** | CCAGAGGTGGACCCTTTGCCTTTC | CACCAGCAGATGCGACACCAGAG |
| ***TBP*** | ACCTTATGCTCAGGGCTTGG | GCCATAAGGCATCATTGGAC |
